# Mechanisms of genomic instability dictate cytosolic DNA composition and dendritic cell mediated anti-tumor immunity

**DOI:** 10.1101/2024.03.08.584184

**Authors:** Shayla R. Mosley, Angie Chen, David N.W. Doell, Siwon Choi, Courtney Mowat, Felix Meier-Stephenson, Vanessa Meier-Stephenson, Kristi Baker

**Affiliations:** Department of Oncology, Cross Cancer Institute, University of Alberta, Canada; Department of Medical Microbiology and Immunology, University of Alberta, Canada

## Abstract

Patients with microsatellite instable (MSI) colorectal cancers (CRC) face better prognosis than those with the more common chromosomal instable (CIN) subtype due to improved anti-tumor immune responses characterized by high cytotoxic T cell infiltration. Previous investigation identified the cytosolic DNA (cyDNA) sensor STING as necessary for chemokine-mediated T cell recruitment in MSI CRCs. Here, we find cyDNA from MSI CRC cells is inherently more capable of inducing STING activation and induces improved cytotoxic T cell activation by dendritic cells (DCs). Sequencing indicates MSI cyDNA is enriched for microsatellites, which upon DC uptake induce anti-tumor immunity in a manner consistent with clinical MSI CRCs. Radiation also modulates cyDNA stimulation capacity through larger cyDNA size and increased mitochondrial DNA content. Identifying highly stimulatory cyDNA arising from genomic instability such as in MSI CRCs allows for optimized development of DNA-based STING agonist therapies to improve responses of CIN CRC patients to immunotherapies.

## Main

Colorectal cancer (CRC) is the second most deadly cancer worldwide, with an estimated 1.9 million people diagnosed per year^1^. The majority of CRCs are initiated by mutations in *APC* and subsequently acquire mutations in *KRAS*, *BRAF*, *PTEN*, *RAD51*, and *TP53* leading to chromosomal instability (CIN)^2–4^. Alternatively, 12-15% of CRCs are established by epigenetic silencing of *MLH1*, resulting in dysfunctional mismatch repair and high levels of point mutations and frameshifts in microsatellite regions of the genome resulting in microsatellite instability (MSI)^5^. Clinically, MSI CRC patients face better prognosis, with an estimated 35% lower risk of death^6^. This can be attributed to their potent anti-tumor immunity characterized by high CD8^+^ cytotoxic T cell infiltration^7^. While this has been linked to high neoantigen production due to the hypermutability of MSI CRCs^5^, this theory is incomplete and fails to explain how MSI tumors in other areas of the body are associated with a worse prognosis than non-MSI tumors^8^. This indicates that neoantigen-independent mechanisms are an important feature of anti-tumor immunity in MSI CRCs. Previously, our lab identified high production of the T cell recruitment chemokines CCL5 and CXCL10 as essential for the increased T cell activation and infiltration of MSI CRCs and showed that their expression was dependent on activation of the cytosolic DNA (cyDNA) sensor STING^9^.

STING is a pattern recognition receptor (PRR) that evolved to recognize microbial DNA in the cytosol^10^. Sensing of cyDNA by cGAS induces its oligomerization and production of the cyclic dinucleotide 2’3’-cGAMP^11,12^. cGAMP then activates STING, which forms a complex with TBK1 and IRF3, each of which are then phosphorylated by TBK1^13^. pIRF3 then translocates to the nucleus to transcribe the type I interferons IFNα and IFNβ, which then induce expression of IFN stimulated genes, such as CCL5, CXCL10, and IRF7, through the JAK/STAT pathway ^10,14^. cGAS can also be activated by endogenous DNA that leaks into the cytosol following DNA damage^14^. This has broad implications for cancers with genomic instability or those treated with DNA-damaging therapies. Indeed, STING has been shown to play a role in cancer prevention^15^ and response to DNA damaging treatments like radiation^16^. As such, STING agonists are currently under clinical trials alone or in combination with checkpoint inhibitors like pembrolizumab^17^.

Dendritic cells (DCs) are key mediators of both anti-tumor immunity and STING dependent immune responses^12^. Exogenous type I IFN leads to DC activation, allowing for co-stimulation and cross-presentation to CD8^+^ T cells^12^. Recently, DCs have been shown to uptake cyDNA and cGAMP from tumor cells by phagocytosis, formation of gap junctions, and uptake of cyDNA-containing exosomes, leading to STING activation and type I IFN production from within the DCs themselves^18,19^.

Although we previously identified STING activation as a neoantigen-independent mechanism of T cell infiltration and activation in MSI CRCs, we observed no significant difference in cyDNA quantity between MSI and CIN cells and proposed there could instead be differences in the nature of the cyDNA produced by these CRCs that underlies the difference in STING activation^9^. Here, we find that early activation of cGAS/STING in DCs by cyDNA arising in tumors with different DNA damage contexts, such as MSI and radiation damage, induce stronger innate and adaptive anti-tumor immunity at equal cyDNA concentration. We identify differences in cyDNA fragment size, subcellular origin, and repetitive sequences found in these highly stimulatory cyDNAs that explain this improved immune activation and show they promote anti-tumor immunity in cold CIN tumors. These findings further support the use of STING agonists for neoadjuvant use in anti-tumor immunotherapy and provide the framework for improved design of DNA based STING agonists.

## Results

### MSI cyDNA more efficiently activates STING in DCs leading to improved anti-tumor immunity

Cross presentation and co-stimulation by DCs is a necessary step for T cell maturation, and has been shown to occur as a result of STING activation^20^. Interestingly, single cell RNA sequencing data from orthotopic MSI and CIN tumors showed an increased type I IFN related gene expression set in DCs from MSI tumors (Fig. 1A, Extended Data Fig. 1A), likely a result of damage associated patterns from the tumor cells. Consistent with this, using MC38 cells rendered MSI or CIN by CRISPR induced mutation^9^, co-culture of BMDCs with MSI or CIN cells labelled by EdU indicated DCs were highly capable of DNA uptake from CRC cells (Extended Data Fig. 1B). Therefore, we aimed to determine whether cyDNA in MSI tumors is inherently more stimulatory to the STING pathway than that of CIN CRCs using cyDNA isolated from MSI and CIN cells. Stimulation of BMDCs with equal cyDNA concentrations showed earlier IFNβ and CXCL10 production by MSI cyDNA than CIN cyDNA (Fig. 1B) despite equivalent uptake (Extended Data Fig. 1C). This early activation was extinguished by later time points when CIN cyDNA shows increased IFNβ and CXCL10 expression (Fig. 1C). MSI cyDNA was also more efficient at inducing IFNβ and CXCL10 expression than herring-testis DNA (HT-DNA), a commonly used STING pathway agonist. This delay in STING activation from CIN cyDNA was not due to greater overall DNA uptake over longer stimulation time frames, as pulsing BMDCs with MSI or CIN cyDNA for only 15 minutes did not alter the kinetics of IFN production (Extended Data Fig. 1D, E). Therefore, our results indicate equivalent amounts of MSI cyDNA lead to quicker and earlier activation of the STING pathway in DCs compared to CIN cyDNA.

**Figure 1.**
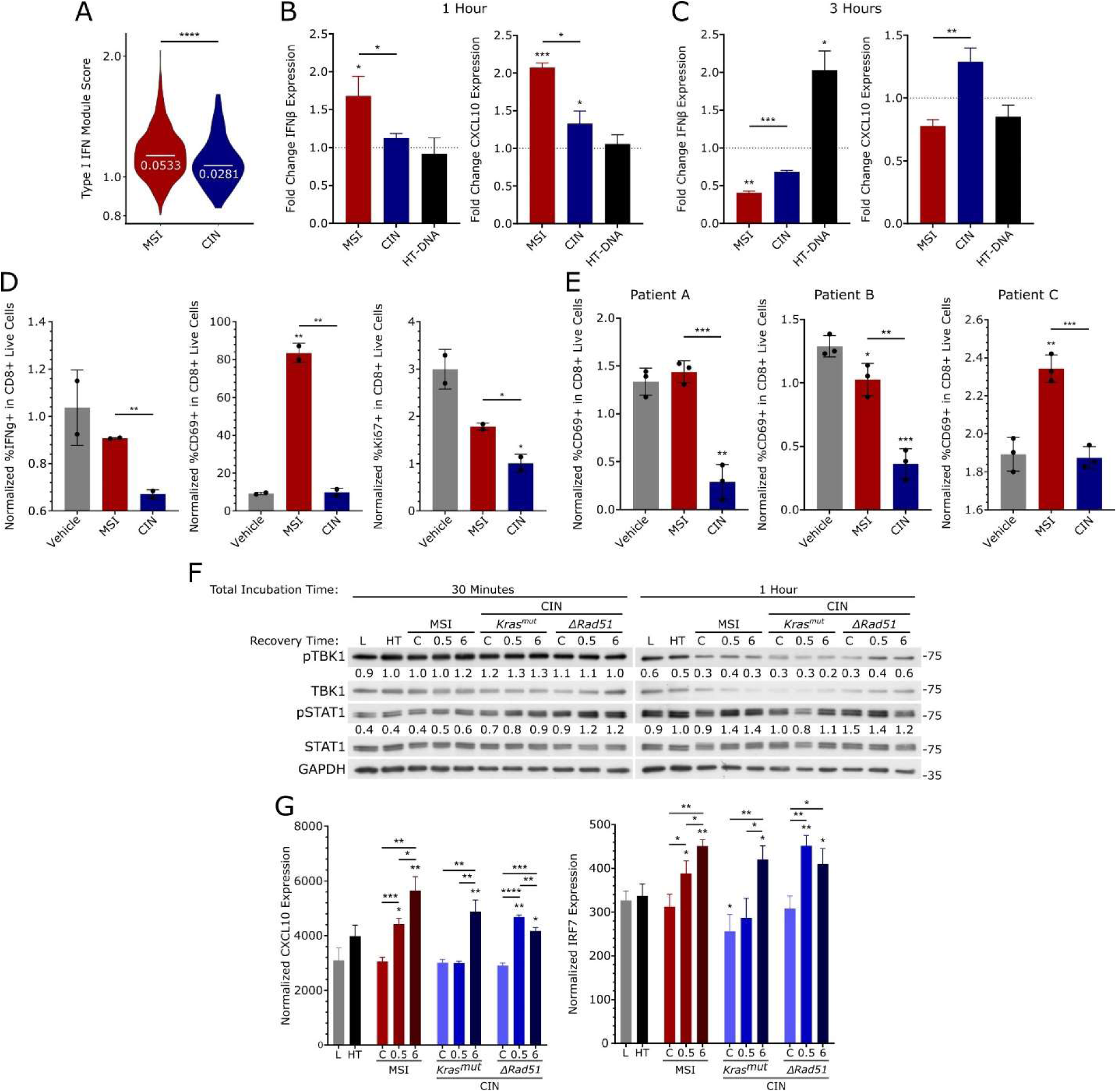
CyDNA from MSI and IR treated cells lead to increased STING signaling in DCs and greater T cell activation. (A) Module score of the Type I IFN gene set expression in DCs from orthotopic MSI and CIN (*Kras^mut^*) tumors analyzed by single cell RNA sequencing. (B, C) 500ng cyDNA isolated from MSI and CIN MC38 cells was used to stimulate BMDCs for 1 (B) or 3 hours (C) before isolation of RNA and qPCR. Data shown as fold change from the lipofectamine vehicle control. n = 3. (D) BMDCs were stimulated with 12.5 or 50ng cyDNA with 0-5µg OVA protein for 30 minutes before removal of stimulus and co-culture with OT-1 CD8^+^ T cells at a 2:1 ratio for 48 hours. T cell activation and proliferation were assessed by flow cytometry. n = 3. (E) CyDNA isolated from *shMLH1* (MSI) or shScramble (CIN) patient organoid cultures was used to stimulate OVA pulsed BMDCs (0-5µg OVA protein) for 30 minutes before removal of stimulus and co-culture with OT-1 CD8^+^ T cells at a 2:1 ratio for 48 hours and examination of T cell activation by flow cytometry. n = 3 (patients A-B) and n = 2 (patient C). (F-G) MC38 cells were treated with 10Gy IR and allowed to recover for 0.5 or 6 hours before isolation of cyDNA. BMDCs were stimulated with 500ng of this cyDNA for 30 minutes, washed to remove the stimulus, and incubated to the indicated times before analysis by western blot (F) or qPCR at 3 hours (G). n = 2 (western blot) and n = 3 (qPCR). L indicates lipofectamine vehicle control, HT indicates HT-DNA control, C indicates cyDNA from non-IR treated cells. Western blot quantifications were determined using ImageJ and normalized to GAPDH loading control. All data shown are from 1 representative replicate. All co-culture data was normalized to T cell only controls. Statistics were determined using unpaired T-tests where significance to the lipofectamine vehicle control is indicated above the bar and significance between experimental samples are above connecting lines, * = p < 0.05, ** = p < 0.01, *** = p < 0.001, **** = p < 0.0001.

Next, we wanted to investigate whether this early STING activation from MSI cyDNA could improve CD8^+^ T cell activation, as this is a key feature of anti-tumor immunity in MSI CRCs^7^. To do this, OVA pulsed BMDCs were stimulated with equal concentrations of MSI and CIN cyDNA and co-cultured with OVA specific OT-1 CD8^+^ T cells. Stimulation with cyDNA from MSI cells led to increased levels of the T cell activation markers IFNγ and CD69, as well as the proliferation marker Ki67 (Fig. 1D). To ensure that this finding was not an artifact of the MC38 CRC cell line, we also isolated cyDNA from control patient derived organoid cultures or matched MSI variants in which *MLH1* was knocked down using shRNA^9^.

Consistent with our previous results, stimulation of BMDCs with *shMLH1* organoid derived cyDNA upregulated more CD69 on CD8^+^ T cells compared to cyDNA from matched controls (Fig. 1E). Therefore, MSI cyDNA leads to improved T cell activation over CIN cyDNA, suggesting that early activation of cGAS/STING in DCs during co-stimulation plays a role in the durable CD8^+^ T cell mediated antitumor immunity in MSI CRCs.

Our data indicates that cyDNA arising from DNA damage due to genetic instability induces differential STING activation. We therefore questioned whether DNA damaging therapies would also alter the stimulatory capacity of cyDNA. To investigate this, we focused on ionizing radiation (IR), a frequently used DNA damaging therapy well known to induce STING activation^13,16^. Therefore, BMDCs were stimulated with cyDNA isolated 0.5 or 6 hours after 10 Gray of IR. Even at equal concentrations, cyDNA from IR treated cells led to increased TBK1 and STAT1 phosphorylation (Fig. 1F), as well as transcription of CXCL10 and IRF7 (Fig. 1G) regardless of whether the treatment occurred in MSI or CIN cells. Therefore, cyDNA from DNA damaging treatments can also influence the stimulation potential of cyDNA, with IR treatment specifically leading to an increased early stimulatory capacity.

### Longer fragments in IR cyDNA lead to increased STING and T cell activation via improved cGAS binding

Previous reports have suggested longer DNA fragments (>50bp) are more capable of inducing cGAS activation^21^. To evaluate the size of cyDNA produced endogenously by MSI vs CIN cells, isolated cyDNA was subjected to gel electrophoresis. This indicated cyDNA in each CRC subtype ranged in size from 10 to 100bp (Extended Data Fig. 2A). To more sensitively investigate cyDNA size differences, we utilized microfluidics based automated electrophoresis with a bioanalyzer (Fig. 2A). Quantification of the resulting curves indicated increased levels of larger cyDNA fragments (>120bp) following IR treatment, suggesting DNA damage from IR led to longer DNA fragments being released into the cytosol. No consistent differences in length were observed between cyDNA from MSI and CIN CRCs.

**Figure 2.**
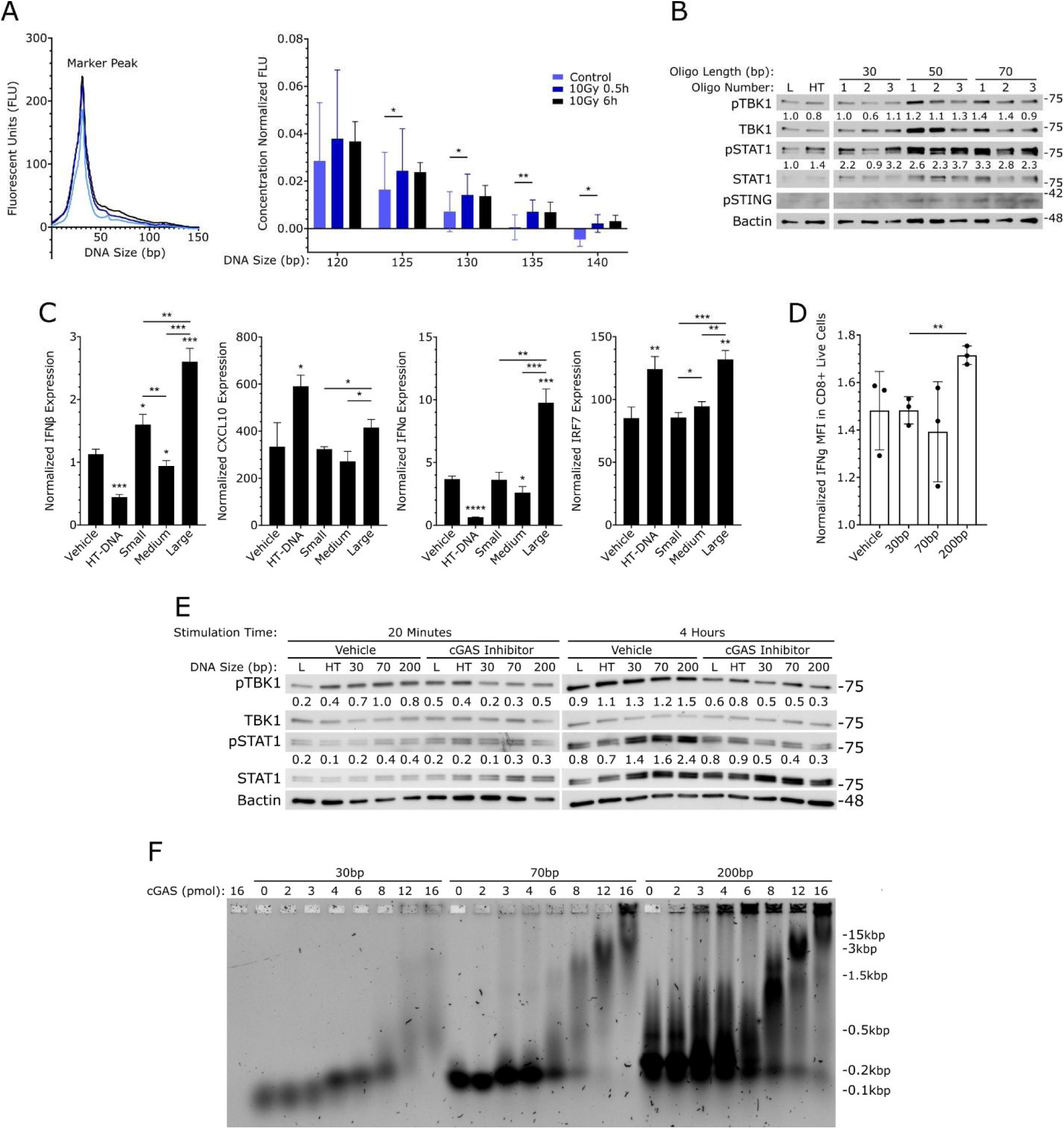
Longer cyDNAs caused by IR treatment induce greater STING pathway activation in DCs. (A) cyDNA isolated from *ΔRad51* CIN cells following IR treatment was run on the Agilent Bioanalyzer 2100 using the High Sensitivity DNA Kit to evaluate cyDNA size. Left: raw data curve with the kit marker peak at 35bp indicated. Right: compiled data of 3 replicates at specific sizes normalized to the quantity of input DNA (determined by Nanodrop). (B) 500ng of oligos of the indicated lengths and scrambled sequence were used to stimulate BMDCs for 3 hours before isolation of protein and western blot analysis. L indicates lipofectamine vehicle control, HT indicates HT-DNA control. Quantification is normalized to βActin loading control and lipofectamine vehicle control. MFI refers to mean fluorescence intensity. (C) cyDNA isolated from MSI cells was separated by size using FPLC to the indicated DNA size range (see Extended Data Fig. 2B). 500ng was used to stimulate BMDCs for 3 (IFNβ, CXCL10, IFNα) or 4 hours (IRF7) before RNA isolation and qPCR. (D) BMDCs were stimulated with 12.5ng oligos of different sizes (oligo #2 of each) for 30 minutes before washing and co-culturing with OT-1 CD8^+^ T cells for 48 hours. T cell activation was evaluated by flow cytometry. Data was normalized to T cell only controls. (E) BMDCs were pre-treated with 3µM RU-521 cGAS inhibitor or DMSO for 18 hours before stimulation with oligos of different sizes (oligo #2 of each) and isolation of protein followed by western blot analysis. Quantifications are normalized to βActin loading control. (F) EMSA of cGAS binding to DNA oligos of different sizes (oligo #2 of each). All data n = 3. Representative replicates shown. Statistics were evaluated using an unpaired T-test. Significance to the lipofectamine vehicle control is indicated above the sample bar, * = p < 0.05, ** = p < 0.01, *** = p < 0.001, **** = p < 0.0001.

To confirm increased cyDNA size induced increased STING activation, oligos of scrambled sequence of 30, 50, or 70bp in length were used to stimulate BMDCs. Consistent with previous reports^21^, increased levels of pTBK1, pSTAT1, and pSTING were observed with 50 and 70bp oligos (Fig. 2B). To determine whether longer cyDNAs produced endogenously also led to this improved activation, MSI cyDNA was separated by size using fast protein liquid chromatography (FPLC) (Extended Data Fig. 2B).

BMDC stimulation with longer cyDNAs also led to increased transcription of IFNα, IFNβ, CXCL10, and IRF7 (Fig. 2C). To investigate whether DNA of increased size could induce stronger cytotoxic T cell activation, BMDCs stimulated with 30, 70, or 200bp oligos were co-cultured with CD8^+^ T cells. Analysis of T cell activation by flow cytometry indicated increased levels of IFNγ following stimulation with larger DNAs (Fig. 2D). This indicates that increased cyDNA size not only leads to stronger STING activation but also to more potent cytotoxic T cell activation by DCs.

Next, we wanted to determine whether this increased activation was dependent on cGAS as other DNA sensors may also stimulate a STING and/or IFN-mediated response^10^. BMDCs were pre-treated with the cGAS inhibitor RU-521^22^ before stimulation with oligos of different sizes. RU-521 strongly decreased pTBK1 and pSTAT1 levels from 70 and 200bp oligo stimulation (Fig. 2E). To determine the underlying mechanism, we then evaluated whether larger cyDNAs exhibited improved binding to cGAS using an electrophoretic mobility shift assay (EMSA), where increased binding is indicated by reduced migration of the complex through an agarose gel. We observed binding of the 70 and 200bp oligos occurred at lower cGAS concentrations than 30bp DNA (Fig. 2F, Extended Data Fig. 2C). Consistent with this, increased cGAS oligomerization, which occurs as cGAS binds to cyDNA^11^, was also observed with the 70 and 200bp DNAs (Extended Data Fig. 2D). Taken altogether, these data suggest longer cyDNA fragments produced by IR treatment lead to stronger cGAS/STING activation in DCs, and thereby stronger T cell activation.

### cyDNA in CRC cells is enriched for specific genetic elements

To begin characterizing cyDNA from MSI vs CIN cells and identify the differences that underly the improved STING activation from MSI and IR induced cyDNA, we performed next generation sequencing on cyDNA isolated from MSI and CIN cells with or without treatment with IR or 5-fluorouracil (5-FU). 5-FU is a common treatment for CRC patients in both FOL-FOX and FOL-FIRI combination therapies and induces DNA damage through integration of modified bases and nucleotide imbalance^2,23^. First, we sought to characterize the overall composition of cyDNA in all samples to understand the nature of cyDNA in general. Although cyDNA mapped to sequences from each chromosome, we observed an unexpectedly high amount of reads from chromosomes 2, 9, 11, and 15 upon comparison to the expected percentage of each chromosome in cyDNA based on base pair size (Fig. 3A). This was combined with unexpectedly low cyDNA contribution from chromosomes 3, 4, 6, 14, 16, 18, and X. CyDNA was also enriched in reads mapping to protein coding gene areas, including both exons and introns (Fig. 3B, C), suggesting selection for euchromatic regions of the genome. Surprisingly, when examining which genes CRC cyDNA mapped to, we observed that 2 of the 5 most common genes across our samples were the immunoglobulin genes *Igk* and *Igh*, which were present in 19.4 and 16.1% of cyDNA samples, respectively (Fig. 3D). This bias for certain chromosomes and genes within cyDNA indicate its composition is not random and suggests there is more DNA damage at these sites overall or that DNA fragments may be exported to the cytosol by a specific mechanism.

**Figure 3.**
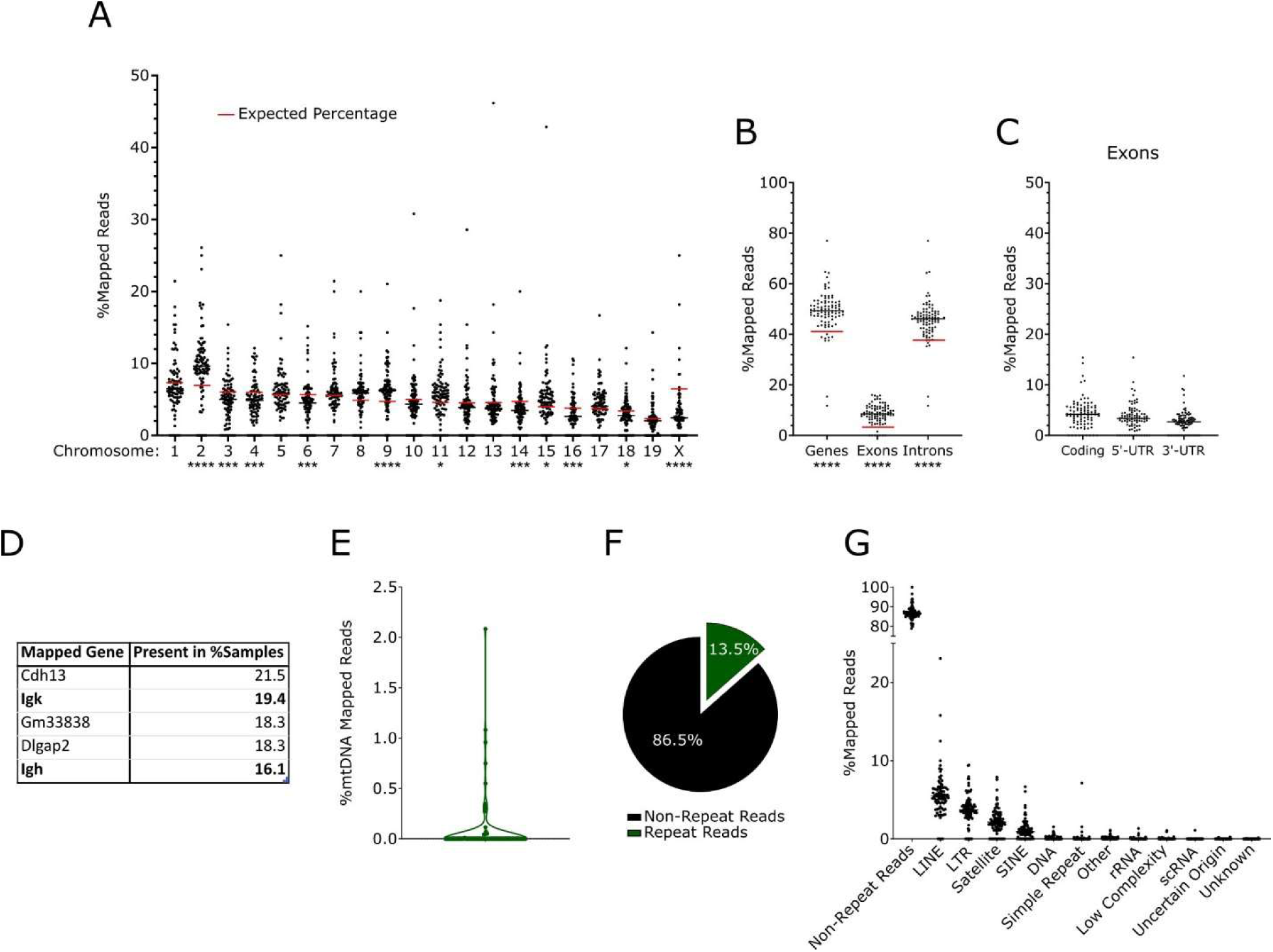
CyDNA contains chromosomal, mitochondrial, and repetitive DNA sequences. cyDNA isolated from MSI and CIN cells with and without IR treatment at 10 Gy (followed by 0.5 or 6 hour recovery) or 5-FU treatment (1.0µM, 1.5µM) for 24 hours was sequenced to examine cyDNA composition. (A) Percentage of cyDNA in each sample from the indicated chromosome. Red line indicates the expected cyDNA contribution of that chromosome based on its size. (B, C) Percentage of cyDNA in each sample mapped to gene coding regions (B), or exon regions (C). Red line indicates the expected percentage of protein coding genes in cyDNA at random. (D) Top 5 most common genes in MC38 cyDNA contains 2 immunoglobulin genes. Ranking determined as percentage of samples the gene was present in. (E) Percentage of reads in each sample that mapped to the mitochondria. (F) RepeatMasker mapping analysis finds an average of 13.5% of reads map to repetitive regions. (G) Repeat types and their average percentage of cyDNA contribution in each sample. Statistical significance compared to the expected percentage (A, B) is indicated under the X axis and were determined using one-sample T-tests, * = p < 0.05, *** = p < 0.001, **** = p < 0.0001.

Many recent studies have investigated how mitochondrial DNA (mtDNA) release into the cytosol affects STING activation^24,25^. We found an overall low amount of mtDNA in the cytosol, comprising an average of 0.08% of cyDNA in CRC cells (Fig. 3E). Another aspect of cyDNA under intense investigation is retrotransposons, such as LINEs and SINEs, which have been shown to play a role in STING activation in senescence, aging, and cancer^26,27^. To evaluate the levels of transposons and other repetitive regions in cyDNA, we examined reads that mapped to regions annotated by RepeatMasker, which identifies repetitive and low complexity regions within the genome, including LINEs, SINEs, and satellites^28^. Using this tool, we found an average of 13.5% of cyDNA mapped to repetitive regions of the genome (Fig. 3F), with the largest contributor being LINEs, LTRs, and satellites (Fig. 3G).

### Genomic instability from MSI and IR produce specific sequence and motif changes within cyDNA

To identify patterns prevalent in highly stimulatory cyDNAs, we investigated our sequencing data for differences between MSI and CIN cyDNA or cyDNA from IR treated or untreated cells. Although mtDNA was a small fraction of cyDNA content (Fig. 3E), we observed increased mtDNA in the cytosol 6 hours after IR treatment, though this difference was not statistically significant (Fig. 4A). In contrast, cyDNA isolated from 5-FU treated cells showed no difference in mtDNA content (Extended Data Fig. 3A). This suggests that IR, but not all DNA damaging agents, promotes the release of mtDNA into the cytosol.

**Figure 4.**
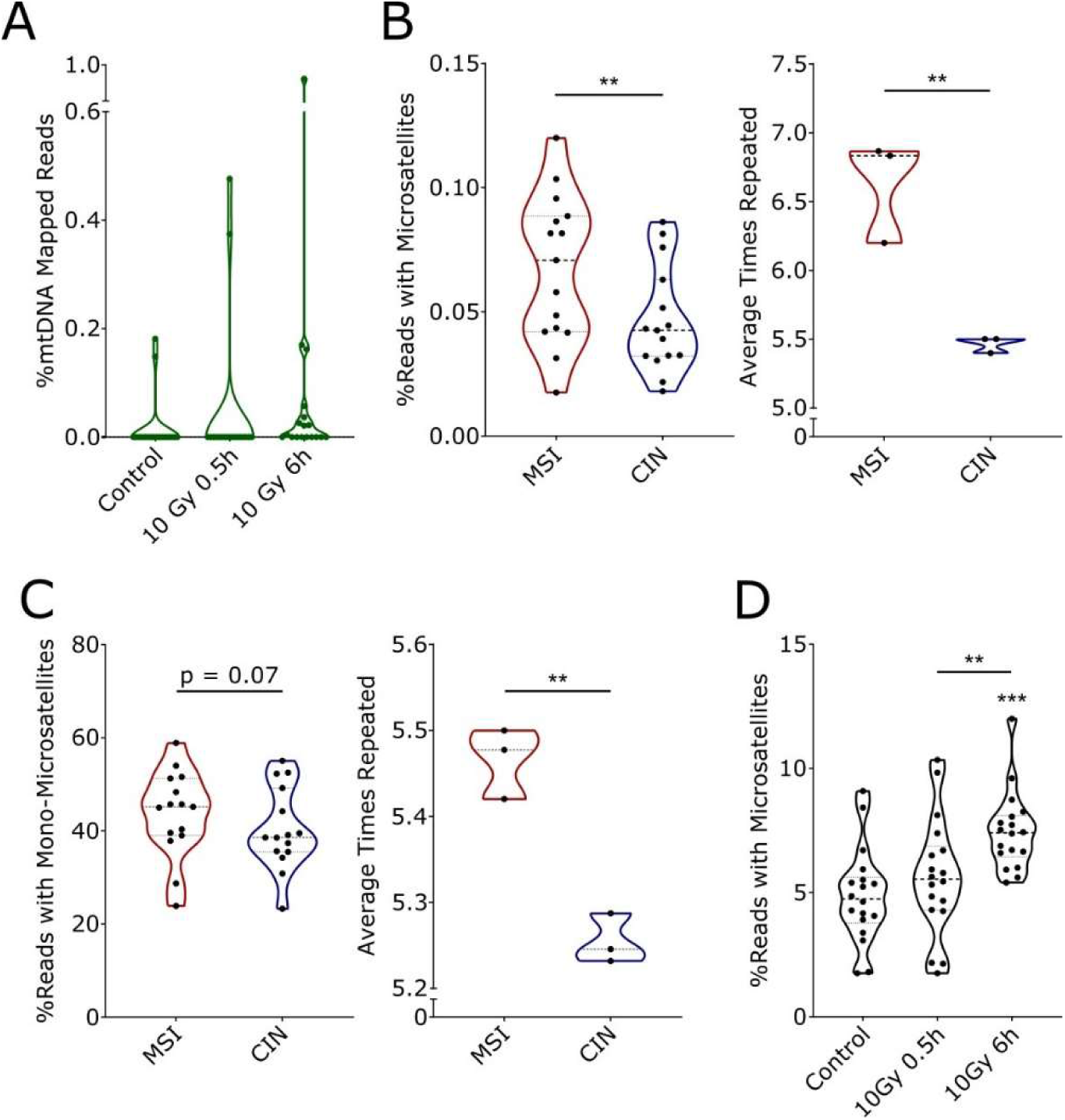
Strongly stimulatory cyDNAs contain higher levels of mtDNA and microsatellites. (A) Percent of cyDNA sequencing reads that mapped to the mitochondria with or without IR treatment (10Gy followed by 0.5 or 6 hour recovery) in all CRC samples. (B) Left: percent of all mapped and unmapped reads that contain >5 2-6bp microsatellite repeats in MSI vs CIN cyDNA for all treatment types combined. Right: number of times each repeat was repeated in untreated cells. (C) Left: percent of all mapped and unmapped reads that contain >5 1bp microsatellite repeats in MSI vs CIN cyDNA from all treatment types combined. Right: number of times each repeat was repeated in untreated cells. (D) Percent of all mapped and unmapped reads that contain >5 2-6bp microsatellite repeats after IR treatment (10Gy followed by 0.5 or 6h recovery) from all CRC subtypes combined. All sequencing was performed with 3 biological replicates and pooled data is shown. Statistics were determined using paired (B left, C left, D) or unpaired (B right, C right) T-tests. Significance to the untreated control is indicated over the sample bar (D), ** = p < 0.01, *** = p < 0.001.

MSI cancers are characterized by point mutations and frameshifts resulting from polymerase slippage, most evident within microsatellite regions^5^, suggesting DNA from damaged microsatellite regions could be more prevalent in the cytosol in these cells. Therefore, we examined the repetitive patterns of cyDNA in our sequencing data. Upon evaluation of RepeatMasker mapped regions in MSI vs CIN cyDNA, we observed differences in the levels of “Other” and SINE annotated regions (Extended Data Fig. 3B). However, no differences were observed in satellite or simple repeat (microsatellite) levels (Extended Data Fig. 3C). We then considered the possibility that highly repetitive regions are well known to be difficult to accurately map, and often require long sequencing reads to do so, which would not be possible in short fragmented cyDNA. Therefore, we performed sequenced based analysis on all cyDNA sequencing reads, regardless of whether they mapped to the mouse genome, using MicRocounter^29^, a Bioconductor package that identifies 2-6 base pair microsatellites in raw sequence data. This revealed increased microsatellite content in MSI cyDNA compared to CIN cyDNA (Fig. 4B). Additionally, microsatellites found in MSI cyDNA were of longer lengths. Similar results were found with single base pair microsatellites (Fig. 4C). Lower GC content in MSI cyDNA was also observed (Extended Data Fig. 3D). Therefore, MSI leads to cyDNA with increased microsatellite repeats.

Surprisingly, increased microsatellites were also observed in the cytosol of all CRCs 6 hours after IR treatment (Fig. 4D), but not after treatment with 5-FU (Extended Data Fig. 3E). Consistent with this, decreased sequence complexity was also observed (Extended Data Fig. 3F). IR and 5-FU both led to changes in cyDNA GC content (Extended Data Fig. 3G). Interestingly, RepeatMasker analysis of IR induced cyDNA also suggested increased levels of DNA transposons in the cytosol after 0.5 hours, which was largely due to the Charlie-hAT Transposon type (Extended Data Fig. 3H, I). To our knowledge, DNA transposons, a type of transposon that translocates via a double stranded DNA intermediate^30^, have not yet been reported within cyDNA. Additionally, the mechanism for their presence in cyDNA is not as clear as for retrotransposons such as LINEs and SINEs, as DNA transposons are traditionally considered fossils, and therefore thought to be no longer capable of translocation in most mammals^30,31^. Given that Charlie-hAT is the most common DNA transposon in humans^31^, understanding the presence of DNA transposons within cyDNA, particularly following IR treatment, could be important in the future for understanding immune responses from IR in CRC patients.

### Cytosolic mtDNA is more stimulatory than cytosolic genomic DNA

Our finding of increased mtDNA in the cytosol following IR (Fig. 4A) was intriguing as the microbial origin of the mitochondria^32^ leads to unique mtDNA properties that may induce improved sensing by cGAS due to its evolutionary selection to sense microbial DNA as non-self. We thus sought to investigate the role of increased mtDNA in the cytosol in cGAS/STING activation. First, to confirm the trending results observed in our sequencing data (Fig. 4A), cyDNA was isolated from MSI and CIN cells following 10 Gy of IR treatment and analyzed by qPCR using mtDNA specific primers. Increased proportions of mtDNA were present in the cytosol 6 hours after IR treatment compared to genomic DNA, specifically in MSI cells (Fig. 5A). To determine whether this increased mtDNA content led to differential cGAS/STING activation, MSI cells were treated with ethidium bromide (EtBr) for 7 days before cyDNA isolation. EtBr has been used extensively in the literature to specifically deplete mtDNA due to its accumulation within the mitochondria and intercalation into highly replicating mtDNA, preventing its replication^33^. After confirming that cyDNA from EtBr treated cells was depleted of mtDNA (Extended Data Fig. 4A), BMDCs were stimulated with equal concentrations of cyDNA from mtDNA depleted or control MSI cells. mtDNA depleted cyDNA induced drastically lower levels of IFNβ and IFNα (Fig. 5B), indicating cytosolic mtDNA is a potent inducer of STING. To understand the relative stimulatory capacity of cytosolic mtDNA compared to genomic DNA, BMDCs were stimulated with differing ratios of mtDNA to genomic DNA isolated from MSI cells and sonicated to equivalent sizes (Extended Data Fig. 4B).

**Figure 5.**
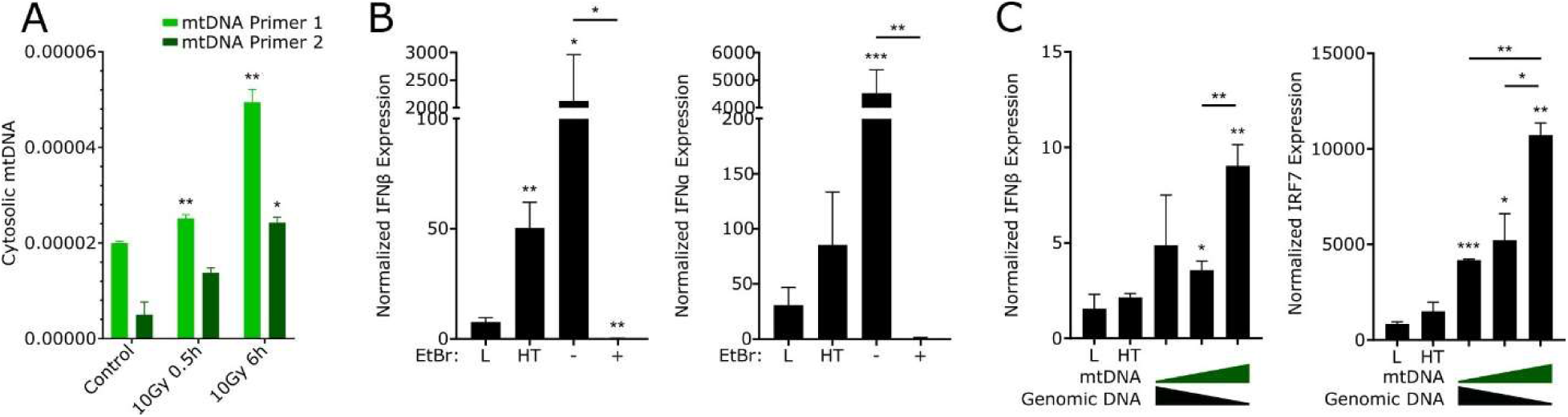
Cytosolic mtDNA induces stronger STING activation in DCs than cytosolic genomic DNA. (A) mtDNA was quantified using qPCR for mtDNA specific sequences in cyDNA isolated from MSI cells treated with 10Gy of radiation and allowed to recover for the indicated time frame. Data shown as proportion of mtDNA in all cyDNA by converting CTs to expression values before normalization to cyDNA quantity determined by Qubit ssDNA Assay Kit. Statistical significance to untreated control is shown over the treated sample bars. (B) MSI cells were treated with EtBr for 7 days to deplete mtDNA before isolation of cyDNA and stimulation of BMDCs for 24 hours. (C) mtDNA and genomic DNA were isolated from MSI cells and sonicated to equal size. BMDCs were stimulated with either mtDNA, genomic DNA, or a 50:50 mix of both for 1 (IFNβ) or 4 (IRF7) hours before isolation of RNA and qPCR. Representative of 3 replicates is shown for all panels. L indicates lipofectamine vehicle control, HT indicates HT-DNA control. Statistical significance was determined by unpaired T-tests. Significance to lipofectamine vehicle control in B and C is indicated over the sample bar, * = p < 0.05, ** = p < 0.01, ** = p < 0.001.

Increased ratios of mtDNA led to higher expression of IFNβ and IRF7 (Fig. 5C), confirming that improved STING activation by cytosolic mtDNA was not an artifact of EtBr use. Taken together, these data indicate that mtDNA is more stimulatory to cGAS/STING than genomic DNA, which may provide an explanation for the improved stimulatory capacity of cyDNA from IR treatment in MSI CRCs. Further investigation will be required to determine what aspects of mtDNA, such as structure, methylation differences, or increased oxidative damage^34–36^, drive this improved immune stimulation.

### Microsatellites in MSI cyDNA lead to improved STING activation through cGAS

Sequencing analysis of MSI and CIN cyDNA indicated more stimulatory MSI cyDNA contained higher levels of microsatellites (Fig. 4B, C). To determine whether microsatellites led to increased STING activation, BMDCs were stimulated with oligos of equal size containing a scrambled sequence or the sequence of a microsatellite containing read identified in our cyDNA sequencing data. As our sequencing data contained reads with microsatellites positioned both in the middle of the cyDNA fragment or on the edge, we positioned the microsatellites in each of these locations in our oligos to determine if this was important. Both edge and middle microsatellite oligos led to increased phosphorylation of TBK1 and STING (Fig. 6A) and increased CXCL10 and IRF7 expression (Fig. 6B) compared to scramble controls, confirming their increased stimulatory capacity. To determine whether endogenous microsatellite containing cyDNA also increased STING activation, microsatellite containing cyDNA was isolated from MSI cyDNA by pulldown of microsatellite motifs found to be enriched in our sequencing data (Fig. 6C, Extended Data Fig. 5A). Stimulation of BMDCs with equal concentrations of this microsatellite-enriched cyDNA or pulldown controls (unmanipulated MSI cyDNA (C), cyDNA with no pulldown step (NP), or scrambled sequence pulldown (S2, S3)) showed increased pTBK1, pSTAT1, and pNF-κB from microsatellite rich cyDNA (Fig. 6D). To examine whether microsatellites could induce the improved T cell activity observed with MSI cyDNA (Fig. 1D, E), BMDCs stimulated with microsatellite or scrambled sequence-containing oligos were co-cultured with CD8^+^ T cells. Microsatellite-containing oligos induced increased levels of Ki67^+^ T cells (Fig. 6E), indicating increased T cell proliferation. Therefore, increased microsatellites in MSI cyDNA leads to improved cGAS/STING activation and cytotoxic T cell activity.

**Figure 6.**
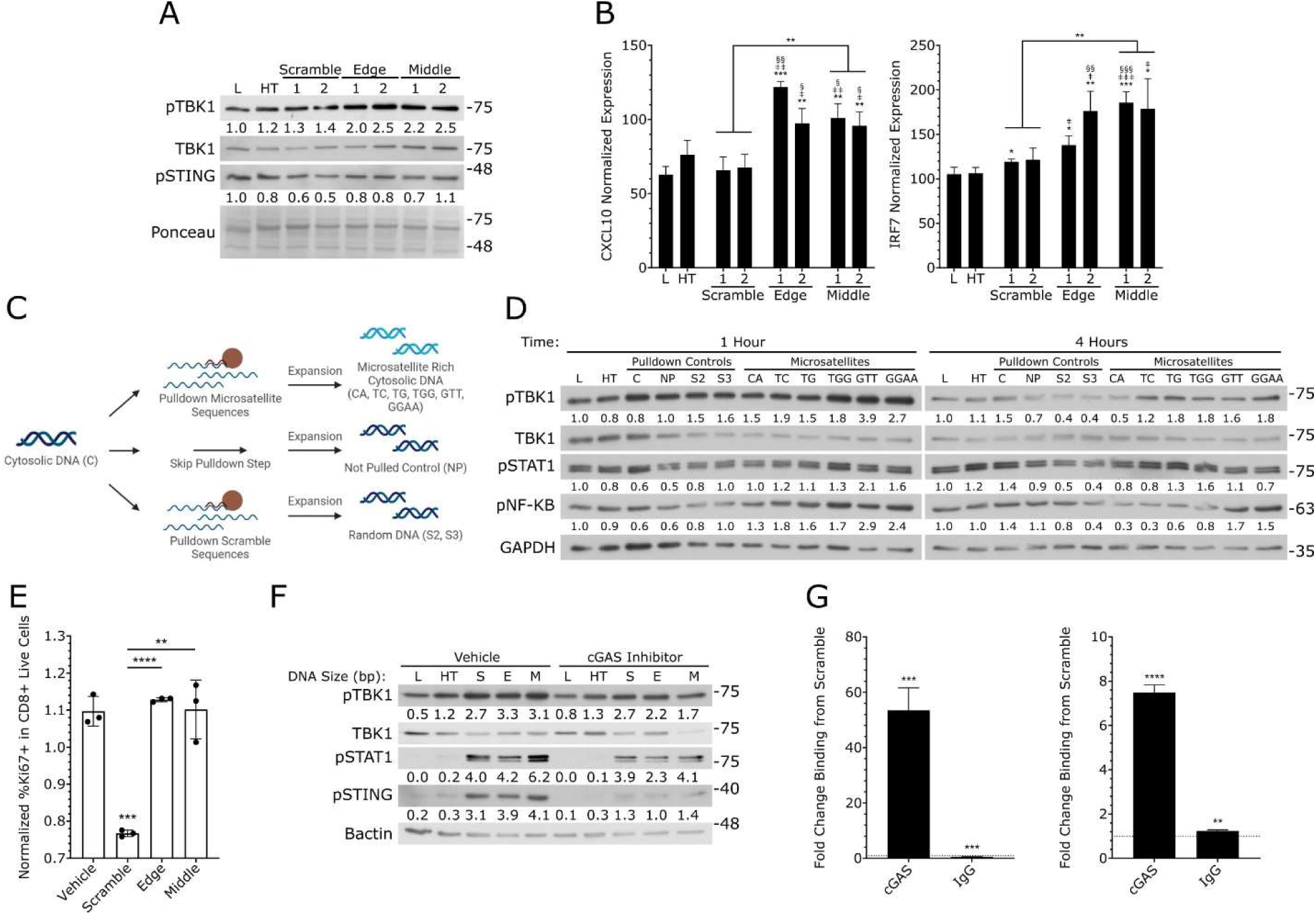
Microsatellite rich sequences induce stronger STING and T cell activation by DCs. (A, B) BMDCs were stimulated with 500ng of oligos containing scrambled or microsatellite sequences on the edge or middle of the oligo before analysis of STING activation by western blot (A) and qPCR (B). Western blot quantifications are normalized to the Ponceau loading control and lipofectamine vehicle control. * indicate statistical significance to the lipofectamine control, ǂ indicates significance to the scramble oligo 1, § indicates significance to the scramble oligo 2. n = 3. (C) Schematic of microsatellite pulldown for isolation of microsatellite rich cyDNAs from cancer cells. cyDNA isolated from MSI cells was pulled down with microsatellite specific or scrambled probes to evaluate the stimulation capacity of microsatellite rich cyDNA. See Extended Data Fig. 5A for more detailed methodology. (D) Microsatellites were pulled down from MSI CRC cyDNA and used to stimulate BMDCs and examine STING activation by western blot. L refers to the lipofectamine vehicle control, HT refers to HT-DNA, C indicates unmanipulated cyDNA, NP indicates the control that skipped the pulldown step but received all other manipulations, S2 and S3 indicate pulldowns with scramble probes, CA, TC, TG, TGG, GTT, GGAA indicate microsatellites pulled down. Quantifications are normalized to GAPDH loading control and lipofectamine vehicle control. n = 3. (E) 12.5ng of oligos containing microsatellite or scrambled sequences (oligo #2 of each) were used to stimulate BMDCs for 30 minutes before washing and addition of OT-1 CD8^+^ T cells for 24 hours. T cell proliferation was then determined using flow cytometry. Data was normalized to T cell only controls. n = 3. (F) BMDCs were pre-treated for 18 hours with 3µM RU-521 cGAS inhibitor before stimulation with 500ng of oligos containing microsatellite or scrambled sequences (oligo #2 of each) for 3 hours and isolation of protein and western blot. S indicates scramble oligo, E indicates the edge microsatellite oligo, M refers to the middle microsatellite oligo. Quantifications were normalized to the GAPDH loading control. n = 2. (G) BMDCs were co-stimulated with 2µg each of scrambled oligo and edge microsatellite oligo, or scrambled oligo and middle microsatellite oligo (oligo #2 of each) for 40 minutes followed by isolation of the cytosolic fraction and ChIP-qPCR of cGAS to evaluate cGAS binding to each oligo. CTs were normalized to the total oligo amount in the cytosol (non-pulled whole cytosolic control) and shown as fold change of microsatellite oligo from scramble oligo. n = 3. Representative replicates shown for all panels. Statistical significance was determined by unpaired T-tests, * = p < 0.05, ** = p < 0.01, *** = p < 0.001, **** = p < 0.0001.

Next, we wanted to investigate whether the improved stimulatory capacity of microsatellite cyDNA was cGAS dependent. Stimulation of BMDCs pre-treated with the cGAS inhibitor RU-521 with microsatellite or scramble oligos showed decreased levels of pTBK1, pSTAT1, and pSTING with cGAS inhibition (Fig. 6F). Consistent with this, BMDC stimulation and protein isolation under non-denaturing conditions indicated increased cGAS oligomerization with microsatellite sequences (Extended Data Fig. 5B). To determine whether increased cGAS activation was due to improved binding to cGAS by microsatellites, BMDCs were competitively co-stimulated with equal concentrations of scrambled and microsatellite containing oligos before pulldown of cGAS and qPCR to examine bound oligo levels. Both edge and middle oligos were more frequently bound to cGAS compared to the scrambled control (Fig. 6G). In contrast, pre-treatment with IN-3, an inhibitor of the PRRs AIM2 and NLRP3 that can also respond to cyDNA^10,34,37^, led to increased pTBK1, pSTAT1, and pSTING from microsatellite containing oligos (Extended Data Fig. 5C), suggesting possible competition for microsatellites in the cytosol between these sensors. Altogether, these data indicate that microsatellites prevalent in MSI cyDNA lead to improved STING activation and T cell activation through increased binding to cGAS.

Traditionally, cGAS has been considered a sequence independent cyDNA sensor^10^, however our data suggests this is not necessarily the case. One possibility is that increased cGAS binding results from secondary DNA structures formed by microsatellite sequences. Indeed, microsatellites are well known to form secondary structures including hairpins, Z-DNA, and G-quadruplexes^38^. Since our microsatellite pulldowns performed in Fig. 6C-D finish with a PCR reaction to convert denatured single stranded cyDNA back to double stranded format for stimulation (Extended Data Fig. 5A), these microsatellite rich cyDNA would likely contain little secondary structure. To determine whether microsatellite cyDNA with increased secondary structure is more stimulatory to STING, this cyDNA was further allowed to form secondary structure by additional denature and re-annealing steps. Stimulation of BMDCs with this highly structured cyDNA led to both earlier and stronger phosphorylation of TBK1 and STING (Extended Data Fig. 5D), confirming that secondary structures formed by microsatellites are a strong activator of cGAS/STING.

### Microsatellite cyDNA induces improved DC and T cell activation in cold CIN tumors

Finally, we wanted to investigate whether this increased STING and T cell activation from microsatellites could improve anti-tumor immune responses in CIN tumors, whose cold immuno-suppressive microenvironment typically drives poor responses to immunotherapies such as checkpoint inhibition^39^. First, BMDCs were stimulated with oligos of 30 or 200bp in size containing scrambled or microsatellite sequences to examine DC activation. Consistent with our previous results, we observed increased levels of pSTING in BMDCs stimulated with the middle microsatellite oligo (Fig. 7A). Increased DC activation was also indicated by increased levels of surface MHCII. Following these stimulations, BMDCs were adoptively transferred by intraperitoneal injection into immunocompetent mice bearing CIN tumors orthotopically implanted into the colon (Fig. 7B). After 2 weeks, CIN tumors were collected and flow cytometry was used to evaluate the anti-tumor immune response (Extended Data Fig. 6A-C). Adoptive transfer of microsatellite oligo stimulated DCs led to increased CD8^+^ T cell infiltration and activation, as shown by the T cell activation markers CD69 and IFNγ (Fig. 7C). This increase was strongest when stimulated with the middle microsatellite oligo and was matched by increased activation of endogenous cross-presenting type I (CD103^Hi^) and high PRR expressing type II (CD11b^Hi^ CD103^Mid^) DCs^40^ in the tumor (Fig. 7D). One concern for STING agonist use is the upregulation of PD-L1 known to occur as a result of increased type I IFN production^13^. However, no increase was observed in PD-L1 expression on CD45^-^ tumor cells upon adoptive transfer of microsatellite stimulated DCs (Extended Data Fig. 6D).

**Figure 7.**
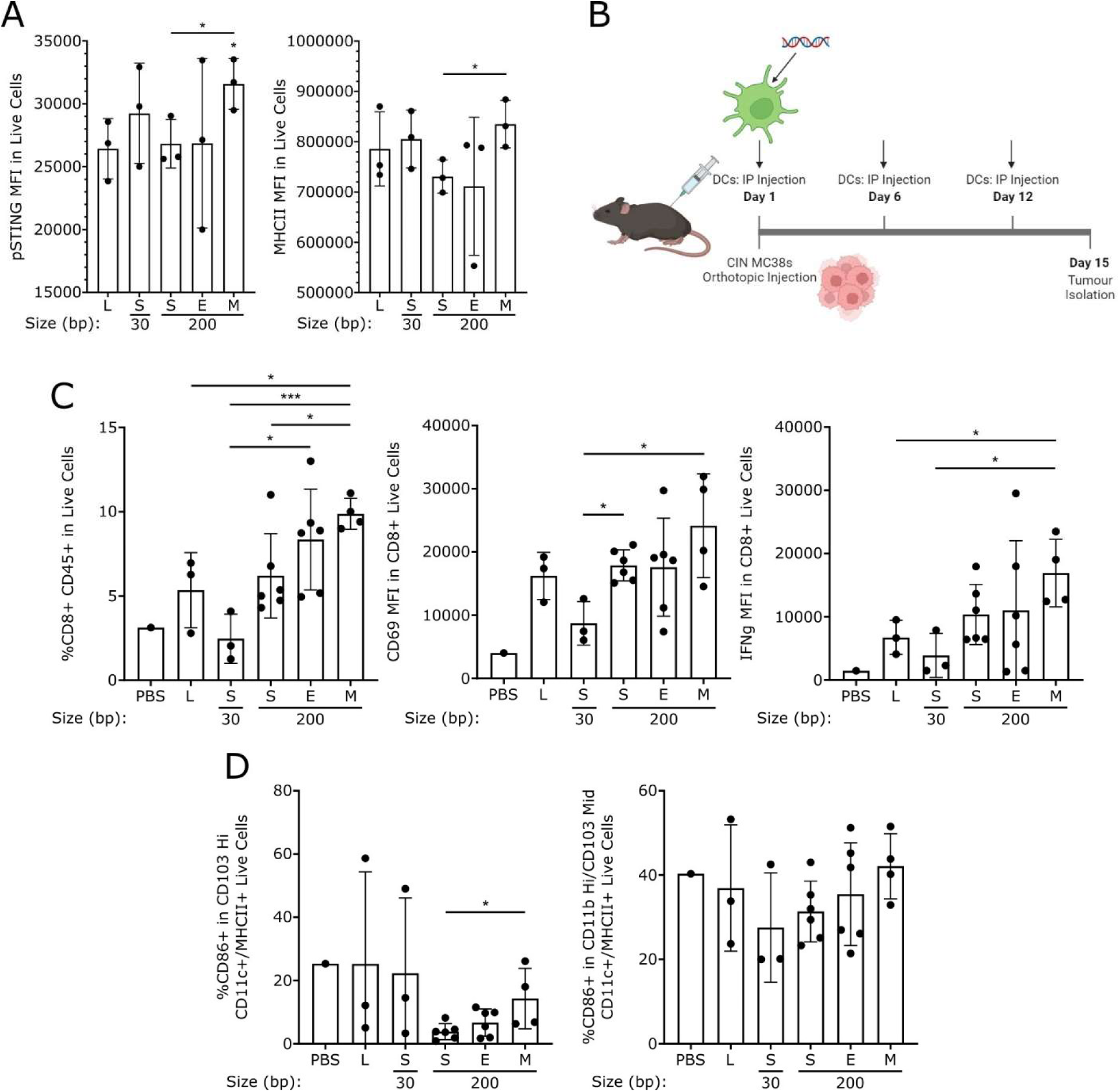
DC activation by microsatellite rich cyDNA leads to improved anti-tumor immunity. (A) BMDCs were stimulated with 30 or 200bp DNA oligos containing scrambled or microsatellite sequences (oligo set 2) for 30 minutes. DC activation and STING signaling were then evaluated by flow cytometry. S indicates scrambled sequence, E indicates edge microsatellite oligo, M indicates middle microsatellite oligo. n =2. (B-D) 2×10^6^ CFDA-stained BMDCs stimulated as in A were intraperitoneally injected into orthotopic CIN tumor-bearing mice at indicated intervals. Tumors were isolated and evaluated for CD8^+^ T cell infiltration and activation (C), and DC activation (D) by flow cytometry. 2-6 mice per experimental group, n = 3 experiments. Representative replicates shown. Statistical significance was determined with unpaired T-tests, * = p < 0.05, *** = p < 0.001.

Therefore, microsatellite cyDNA leads to improved overall anti-tumor immunity in the form of increased T cell activation and infiltration, consistent with the clinical picture of MSI CRCs. Induction of the STING pathway in this manner strengthens the anti-tumor immune response in cold CIN tumors and may thus help to sensitize CIN tumors to immunotherapy treatments for overall improvement of CIN patient prognosis.

## Discussion

Uncovering the mechanisms underlying successful anti-tumor immunity in MSI CRCs provides the opportunity to identify novel treatment strategies that are effective in immunologically cold tumors. One such neoantigen-independent mechanism is activation of cGAS/STING and type I IFN signalling pathways, which we have previously shown are important for MSI CRC anti-tumor immunity. However, our previous work and that of others did not identify the underlying mechanism for activation of these pathways in MSI CRCs. Here, we have identified the specific stimulatory characteristics of cyDNA in MSI CRCs that activate these pathways and further show that this can be replicated using DNA damaging radiotherapy. Specifically, we observed enrichment of sequence patterns in cyDNA of genetically instable tumors that leads to highly potent activation of cGAS/STING. These patterns include microsatellites in MSI cyDNA that we show lead to increased cGAS binding and STING pathway activation in DCs and translate to improved T cell-mediated antitumor immunity in CIN CRCs *in vivo*. We further show that induction of DNA damage by IR treatment leads to larger cyDNA fragment sizes and increased mtDNA leakage into the cytosol, both of which potently increase STING and T cell activation. Notably, we have also identified DCs in the tumor microenvironment as key drivers of cGAS/STING-mediated anti-tumor immunity in MSI CRCs in response to tumor-derived cyDNA.

We are not the first to identify STING as a key player in converting immuno-suppressive cold tumors to hot, with STING agonists currently in clinical trials in combination with checkpoint inhibition therapies^17^. Indeed, Guan *et al*^41^ observed that cyDNA production in MSI cancers could occur from hyperexcision of DNA by EXO1 due to loss of *MLH1* expression. Further evaluation will be needed to determine if EXO1 plays a role in the increased microsatellite presence in MSI CRC cyDNA. A recent study by Wang *et al*^42^ also identified STING activation in type I DCs, but not tumor cells, as necessary for effective CD8^+^ T cell mediated tumor rejection by STING agonists, consistent with our finding of DC mediated anti-tumor immunity resulting from MSI cyDNA. Our work also confirms previous findings in the literature that DNA fragment sizes >50bp led to improved cGAS activation^21^. This is thought to be due to the improved ability of cGAS to oligomerize, and therefore produce 2’3’-cGAMP, on longer DNA strands^43^. Additionally, we determined that endogenous cyDNA contains different sizes, and treatments like IR can increase the proportion of large cyDNA fragments. Our work can thus serve as a foundation for improving the design of STING agonist-based neoadjuvant therapies.

We performed next generation sequencing on cyDNA from cancer cells with different sources of genomic instability and DNA damage to characterize the resulting changes in composition. To our knowledge, we are the first to sequence cyDNA in this manner. This sequencing-based screen was chosen due to the sequence-specific nature of MSI associated damage and therefore its resulting cyDNA. This strategy allowed us to identify microsatellites as being more prevalent in MSI cyDNA and having a greater ability to induce STING activation through stronger cGAS binding leading to overall improved anti-tumor immunity. This sequence dependent mechanism contradicts the current dogma of cGAS as a sequence independent cyDNA sensor^10^. One possibility is that microsatellite sequences could form secondary DNA structures that improve cGAS binding and oligomerization around the structured cyDNA strand. Consistent with this, Herzner *et al*^44^ found that DNAs as small as 12bp were capable of inducing strong IFN induction with the addition of Y-form ends. We also observed increasing secondary structure by re-annealing our microsatellite containing cyDNA led to substantially increased STING activation.

However, microsatellite rich cyDNA of a largely linear conformation also led to increased STING activation, suggesting secondary structure does not completely explain improved immune responses by microsatellites. Other sources have also identified sequence dependent variations in STING activation.

Herzner *et al*^44^ observed that activation by small Y-form DNA required guanosine ends, while Gentili *et al*^45^ found improved IFN production from AATGG centromeric repeats at DNA sizes of 20-30bp. These data suggest that the traditional sequence independent view of cGAS may not be absolute and requires further study.

An important limitation of our work is that our sequencing method did not capture the many other variations within cyDNA that are likely present, such as chemical modifications, methylation, structure, and epigenetic changes. Indeed, a study by Balzarolo *et al*^46^ showed increased IFNβ expression and DC activation upon stimulation using m6A methylated DNA oligos compared to unmethylated controls. This modification is rare in mouse and humans but common in prokaryotes. As m6A is enriched in mtDNA^35^, this finding may have important implications for the increased STING activation by cytosolic mtDNA. An additional limitation is that this study focused on analysis of free fragmented cyDNA and the role of cyDNA arising from micronuclei was not investigated. Interestingly, a recent study by MacDonald *et al*^47^ found the epigenetic marker H3K79me2 in micronuclei led to increased cGAS co-localization and IFN stimulated gene expression, suggesting that epigenetic and protein based differences in micronuclei may affect the level of STING activation.

Overall, this work identifies novel patterns of cyDNA composition arising from different forms of genomic instability that activate STING in DCs and lead to improved T cell infiltration and activation.

Moving forward, this will enable improved design of DNA-based STING agonists as a neoantigen-independent immunotherapy and sets the stage for further analysis of how such agonists can be crafted to maximally activate STING-mediated anti-tumor immunity without also triggering tumor-promoting inflammation.

## Methods

### Cells

To model MSI and CIN CRCs, we previously developed CRISPR mutations in the MC38 mouse CRC cell line^9^. MSI is modeled by deletion of *Mlh1*, a gene silenced in the majority of sporadic MSI CRCs^6^. CIN is modeled by an activating Kras mutation (*Kras^mut^*), as is present in 40% of CRCs^2^ or deletion of *Rad51*, a gene with expression at low or undetectable levels in 88% of CRCs^4^. Mutated cells were cultured continuously for 6 months before use to allow accumulation of mutations consistent with each CRC subtype. Cells were cultured in DMEM (10% FBS, 1% Penicillin/Streptomycin, 1% HEPES) at 37°C with 5% CO2.

BMDCs were generated as previously described^48^. Femur and tibia bones were harvested from C57BL/6 mice and bone marrow was flushed out with PBS. After a PBS wash, cells were plated in non-tissue culture treated plates at 5×10^6^ cells/8mL BMDC growth media (RPMI, 10% supernatant from B16-GMCSF cells, 5% FBS, 10mM HEPES, 1% Penicillin/Streptomycin, 50uM β-Mercaptoethanol). 8mL fresh media was added at days 3 and 5, and non-adherent cells were collected and frozen down for use on day 7.

For T cell isolation, the spleen and lymph nodes were collected from OT-1 C57BL/6 mice before negative selection of CD8^+^ T cells using the EasySep Mouse CD8^+^ T Cell Isolation Kit (StemCell) as per manufacturer instructions.

### Human CRC Organoids

Human CRC organoids were generated from resected tumors as described previously^9,49^. Briefly, tumors were dissociated for 1 hour at 37°C in DMEM with 2.5% FBS, 75 U/ml Collagenase XI (SigmaAldrich), 125 µg/ml Dispase II (SigmaAldrich). Following filtration, cells were plated at 500-1000 per well in growth factor reduced Matrigel (Corning) and cultured in basal crypt media (Advanced DMEM/F12 containing 10% FBS, 2 mM Glutamine, 10 mM HEPES, 1 mM N-Acetylcystein, 1X N2 supplement, 1X B27 supplement, 10 mM Nicotinamide, 500 nM A83-01, 10 µM SB202190, 50 ng/ml EGF) (ThermoFisher) mixed 1:1 with conditioned supernatant from L-cells expressing Wnt3a, R-Spondin and Noggin (ATCC #CRL-3276)^50^. All work with human samples was approved by the Health Research Ethics Board of Alberta Cancer Committee and carried out after obtaining informed patient consent.

Primary MSI variants of the organoids were generated previously using lentiviral transduction with the pLKO.1 system^9,51–53^. Briefly, organoids were pretreated for 4-5 days with 10 mM Nicotinamide, dislodged from the plate by pipetting, and treated for 5 minutes at 37°C with TrypLE Express (Life Technologies). Organoids were mixed with concentrated lentivirus along with 8 µg/ml Polybrene and 10 µM Y27632 (SigmaAldrich) and seeded into a 96-well plate. The plate was centrifuged for 60 minutes at 600g at 32°C and then incubated at 37°C for 6 hours. The organoids were then embedded in Matrigel and cultured in media containing 50-100 µg/ml Hygromycin to select for successful transduction. Gene knockdown was verified by western blot.

### CyDNA Isolation

CyDNA was isolated as previously described^54^. Briefly, 5×10^6^ MC38 cells were collected by trypsinization before extraction of the cytosolic fraction by incubation in cytosolic extraction buffer (150mM NaCl, 50mM HEPES, 200µg/mL Digitonin, 1M Hexylene Glycol) for 10 minutes on ice and centrifugation at 2000g for 10 minutes at 4°C. The supernatant was collected and cytosolic protein and RNA was then removed by Proteinase K (1mg/mL at 55°C for 1 hour, Fisher) and RNase A (500µg/mL at 37°C for 1 hour, Invitrogen) treatment, each of which were followed by phenol/chloroform/isoamyl alcohol extractions. To isolate cyDNA from patient organoid cultures, 24 confluent wells of organoids were collected using 0.5mL/well Cell Recovery Solution (Corning) on ice. After spinning down at 500g for 5 minutes at 4°C, cells were resuspended in 3mL TripLE Express (Gibco) and incubated for 5 minutes at room temperature. Following PBS wash, cells were resuspended in a modified cytosolic extraction buffer (150mM NaCl, 50mM HEPES, 100µg/mL Digitonin, 1M Hexylene Glycol), and the remainder of the cyDNA isolation procedure outlined above was performed. Confirmation of undetectable mitochondrial and nuclear contamination was performed by western blot of the cytosolic and pelleted fractions before protein removal.

### scRNASeq

scRNAseq was previously published by us on orthotopically grown MSI and CIN CRCs and deposited as dataset GSE178706 at the NCBI Gene Expression Omnibus^9^. Gene signature expression analysis (GSEA) was performed as in the original publication to identify Gene Ontology (GO) signatures associated with each CRC subtype^55^.

### BMDC Uptake of Endogenous CRC DNA

DNA in the CRC cells was labeled with the Click-iT EdU Cell Proliferation Kit (ThermoFisher). CRC cells were then cocultured at a 1:1 ratio with BMDCs for 4 hours and analyzed for uptake of CRC DNA by flow cytometry on the CytoFlex S cytometer (Beckman Coulter). Analysis was performed in FlowJo.

### BMDC CyDNA Stimulations

BMDCs were thawed at 37°C for 3 minutes and resuspended in RPMI (10% FBS, 1% HEPES, 1% Penicillin/Streptomycin, 50µM β-Mercaptoethanol) before plating 0.5 x 10^6^ cells/2mL in 24 well plates. BMDCs were then stimulated using 500-1000ng DNA as indicated and 0.125µL/mL Lipofectamine 2000 (Invitrogen) as per manufacturer instructions. For pulse stimulations (Fig. 1F, G, Extended Data Fig. 1D, E), BMDCs were stimulated in 2mL tubes and stimuli containing media was removed after 15 or 30 minutes as indicated before PBS wash and plating. DNA oligos are listed in Supplemental Table 1 and were annealed by combining 20µM of each oligo, 10µL NEB Buffer 2, and up to 100µL nuclease free H2O before incubation at 95°C for 4 minutes, 70°C for 10 minutes, and slow cooling to room temperature overnight. PicoGreen staining of cyDNA (Extended Data Fig. 1C) was performed at 37°C for 2 hours at 8µL/mL. In stimulations utilizing RU-521, BMDCs were pretreated for 18 hours and throughout the stimulation at 3µM. For IN-3, BMDCs were pretreated for 0.5 hours and throughout the stimulation at 10µM. For isolation of protein, PBS washed BMDCs were lysed in lysis buffer (50mM Tris-HCl pH 7.5, 150mM NaCl, 50mM Sodium Pyrophosphate, 1mM EDTA, 0.5% NP40, 1% Triton X-100, 2mM Sodium Orthovanadate, 1% Protease Inhibitor (Sigma)) and incubated for 30 minutes at 4°C on a rotisserie and centrifugation at 18300g for 15 minutes at 4°C to pellet and remove lipids.

Protein concentration was evaluated via the Pierce BCA Protein Assay Kit and 2-10µg of protein was loaded onto SDS-PAGE gels for western blot analysis. Antibodies are listed in Supplemental Table 2.

To assess gene expression, RNA was isolated using TRIzol (Invitrogen) and reverse transcribed with the MultiScribe Reverse Transcriptase Kit (Applied Biosystems). qPCR reactions were performed using the QuantStudio 6 Real-Time PCR System (Applied Biosystems) with the primers listed in Supplemental Table 3 and the PowerUp SYBR Green Master Mix (Applied Biosystems). All data shown is normalized to GAPDH unless otherwise stated. All graphs were made in GraphPad Prism 8 and all figures were organized in Inkscape.

### cGAS Pulldown

For examination of microsatellite binding to cGAS, Protein G Dynabeads (Invitrogen) were washed 3 times in blocking solution (PBS, 0.5% BSA) and incubated with 0.4µg of cGAS or IgG antibodies listed in Supplemental Table 2 overnight at 4°C on a rotisserie. Beads were then washed 3 times in blocking solution. 2×10^7^ BMDCs were plated in 6 well plates before stimulation with 2µg DNA and 0.125µL/mL Lipofectamine 2000 (Invitrogen) as described by the manufacturer. Cross links were formed by addition of formaldehyde to the media for a final concentration of 1%, incubated for 10 minutes, and quenched using glycine at a final concentration of 0.125M. The cytosolic fraction was then isolated by resuspension of cells in cytosolic extraction buffer and incubation on ice for 15 minutes before centrifugation for 10 minutes at 2000g at 4°C. The cytosolic fraction in the supernatant was then collected. An aliquot was removed for the whole cytosolic fraction control and the remaining lysate combined with the beads. This mixture was then incubated overnight at 4°C on a rotisserie. Beads were washed 5 times with wash buffer 1 (50mM HEPES, 500mM LiCl, 1mM EDTA, 1% NP40, 0.7% Na-Deoxycholate) and twice with wash buffer 2 (TE, 50mM NaCl). Protein complexes were eluted (50mM Tris-HCl pH 8, 10mM EDTA, 1% SDS) by incubation for 1 hour at 65°C on a shaking heat block. Cross links were reversed from pulldown and whole cytosol samples by overnight incubation at 65°C. After addition of TE, samples were incubated with RNase A (Invitrogen) at 0.2mg/mL for 2 hours at 37°C, then Proteinase K (Fisher) at 0.2mg/mL for 30 minutes at 55°C before phenol/chloroform/isoamyl alcohol extraction.

### BMDC and T-cell Co-cultures

5×10^4^ BMDCs were plated in 96 well plates in 50µL RPMI (10% FBS, 1% HEPES, 1% Penicillin/Streptomycin, 50µM β-Mercaptoethanol) and stimulated with 12.5-50ng DNA and 0.125uL/mL Lipofectamine 2000 (Invitrogen) as per manufacturer instructions and 0-5µg/mL OVA protein (Millipore Sigma) for 30 minutes. BMDCs were then washed 3 times with warm PBS before addition of 1×10^5^ OT1 CD8^+^ T cells (2:1 T cell to BMDC ratio) and incubation at 37°C for 24-48 hours. Cells were restimulated for 4 hours with 0.5µg/mL PMA, 50µg/mL Ionomycin, and 2 hours 2µM Monensin before flow cytometry using the antibodies listed in Supplemental Table 2, Zombie Aqua Fixable Viability Kit (BioLegend), and the FOXP3/Transcription Factor Staining Kit (eBioscience) and run on the CytoFlex S cytometer (Beckman Coulter). Analysis was performed in FlowJo.

### CyDNA Separation by FPLC

CyDNA samples were separated by size exclusion chromatography using a HiLoad Superdex 75 10/300 column (Cytiva) pre-equilibrated with FPLC buffer (20mM HEPES, pH 7.5, 100mM KCl, 1mM EDTA). With each run, 1 mL of sample was loaded onto the column and eluted at 0.65 mL/min with FPLC buffer. Peak fractions were pooled and concentrated using a 3 kDa spin column (Cytiva) and used in BMDC stimulations.

### cGAS Oligomerization

Following BMDC stimulation as outlined above, protein was instead collected in non-denaturing lysis buffer (50mM HEPES, 150mM NaCl, 10% glycerol, 2mM EDTA, 0.5% Triton X-100) before addition of 5X non-denaturing loading dye (0.25M Tris-HCl pH6.8, 30% glycerol, 0.25% bromophenol blue, 1% deoxycholate). 2-10µg of protein was run on 4-15% BioRad Mini-PROTEAN TGX Gels pre-run at 40 mA for 30 minutes in non-denaturing running buffer (0.025M Tris-HCl, 0.192M glycine) with 0.2% deoxycholate in the cathode chamber before performing western blot analysis.

### Electrophoretic Mobility Shift Assay (EMSA)

EMSA was performed as described in^43^. 1pmol of DNA oligo was combined with the indicated concentration of recombinant human cGAS protein (Cayman Chemicals) in EMSA buffer (150mM NaCl, 20mM HEPES) for a final volume of 10µL before incubation for 30 minutes at room temperature. On ice, 1µL of 10X BlueJuice Gel Loading Buffer (Invitrogen) was added before loading onto a 2% agarose gel at 4°C. Gels were then run at 4°C in 0.5X TBE before staining in 1/10000 SYBR Gold (ThermoFisher Scientific) for 1 hour at room temperature on a rocker and visualization on the Typhoon 9400 Variable Mode Imager using the A^488^ channel.

### Sequencing

CyDNA isolated from MSI and CIN MC38 cells following γ-IR or 5-FU (Sigma-Aldrich) treatment was sequenced by Lucigen using the Mini-Y Adapter Library kit specialized for small DNA strands. Genomic mapping to mouse GRCm38 and RepeatMasker analysis was performed by Lucigen. Read quality was evaluated by Python Bio.SeqIO^56^ and FastQC^57^. Rsamtools^58^ in Bioconductor was used for BAM file analysis of chromosome mapping. NCBI RefSeq gene annotations were downloaded from UCSC and compared to mapping data using rtracklayer^59^ and GenomicRanges^60^ in Bioconductor.

Expected gene, exon, and intron coverage in the genome was taken from Yue *et al*^61^. Before analysis of unmapped reads, reads identified as non-mouse with BLASTn (Database: Nucleotide collection (nr/nt), Organism: exclude *Mus musculus*, Expect Threshold: 0.01, Optimize for: Highly similar sequences (megablast), Percent Identity Filter >99%, Query Alignment Start Filter 1-10) were removed to filter out contaminating bacterial and vector reads from the kit using Python. Filtering by this method removed 96.8% of contaminating reads. Filtered reads were then paired using BBMerge^62^ on default settings.

Therefore, unmapped read analysis was performed on all reads that BLAST identified as mouse or failed to align with the high BLASTn thresholds listed above. GC content and local composition complexity was evaluated using Bio.SeqUtils and BioSeqIO in Python^56^. 2-6 bpmicrosatellites were identified by MicRocounter^29^ and had to repeat a minimum of 5 times to be counted. 1bp microsatellites were identified using Python and Bio.SeqIO by looping through each base in each read and determining if the following base matched. If the following >4 bases were the same as the current base iteration, the sequence was considered a mono-base microsatellite and these bases were skipped before the loop continued. IR and 5-FU graphs with data combined from all subtypes contain 2 clones of MSI cells (*ΔMlh1*), 4 CIN cell types (2 *Kras^mut^* clones, *ΔRad51*, unmutated MC38 CRISPR vehicle control), and *Polε* deficient (*ΔPolε*).

### MtDNA Depletion and Isolation

To deplete mtDNA, MSI and CIN cells were treated with 150ng/mL EtBr for 7 days before cyDNA isolation. mtDNA was isolated using the Mitochondrial DNA Isolation Kit (Abcam) followed by the Qiaprep Spin Miniprep Kit (Qiagen). Genomic DNA was isolated via the GeneJet Genomic DNA Purification Kit (Thermo Scientific). Sonication was performed using the Bioruptor Pico (Diagenode) with 25 cycles of 30 seconds on, 30 seconds off.

### Microsatellite CyDNA Pulldown

Pulldown protocol was modified from^63^. 5µg MSI cyDNA was blunted and terminal phosphates confirmed using the NEB Quick Blunting Kit, before ligation to microsatellite expansion adapters containing an EcoRV restriction site (listed in Supplemental Table 1) using the T4 DNA Ligase Kit (NEB) at a 5:1 ratio overnight at 16°C. Single stranded adapters were previously annealed to double stranded by combination of 20µM of each oligo, 10uL NEB Buffer 2, and nuclease free H2O to 100µL and incubation at 95°C for 4 minutes, 70°C for 10 minutes, and slow cooling to room temperature overnight. After purification of DNA by phenol/chloroform/isoamyl alcohol extraction, microsatellites were pulled down using biotinylated microsatellite probes (containing 5 repeats) listed in Supplemental Table 1 as follows. EasySep Streptavidin magnetic beads (StemCell) were washed in 100µL 6X SSC (For 20X SSC: 3M NaCl, 0.3M sodium citrate) 3 times before incubation overnight on a rotisserie at 4°C in blocking buffer (2% BSA, 0.5% SDS, PBS). After another 3 washes in 100µL 6X SSC, beads were resuspended in 100uL 6X SSC. Purified DNA was then hybridized with the biotinylated DNA probes by combining 10µg DNA, 500pmol probe, 60µL 6X SSC, and nuclease free H2O to 100µL. This mixture was heated to 95°C for 10 minutes and then incubated at hybridization temperature (approximately 8°C below probe Tm) for the specific probe used (CA, TC, TG: 15°C, TGG: 53°C, GTT: 34°C, GGAA: 55°C).

Following this incubation, samples were always kept on ice or at 4°C. The DNA hybridization mixture was then combined with the beads and incubated on the rotisserie for at least 20 minutes at 4°C. The hybridization mixture was then removed from the beads before bead washing on the rotisserie for 5 minutes with 200µL 3X SSC twice, 200µL 2X SSC twice, 200µL 1X SSC twice, and resuspended in 30µL nuclease free H2O. Beads were then incubated at 70°C for 10 minutes on a shaking heat block and the supernatant containing the pulled down microsatellite cyDNA was collected. As this collected DNA is single stranded, PCR was performed using the Q5 High-Fidelity DNA Polymerase Kit (NEB) and primers specific for the microsatellite expansion adapters (1X Q5 Reaction Buffer, 200µM dNTPs, 0.4µM each primer, 100ng template, 0.02U/µL Q5 High-Fidelity DNA Polymerase) (Cycling conditions: 98°C for 5 minutes, 35 cycles of: 98°C for 30 seconds, 69°C for 30 seconds, and 72°C for 30 seconds, and final extension at 72°C for 2 minutes). Adapter sequences were then removed by restriction digest (PCR reaction product, 1X rCutSmart NEB Buffer, 5 U EcoRV-HF (NEB)) at 37°C for 1 hour, then 20 minutes at 65°C. Phenol/chloroform/isoamyl alcohol extraction was performed for final DNA purification. To re-anneal microsatellites for secondary structure formation (Extended Data Fig. 5D), following the PCR step DNA was heated to 95°C for 4 minutes, 70°C for 10 minutes, then allowed to slowly cool to room temperature overnight before continuation of the isolation. Experimental schematics in Fig. 6C, 7B and Extended Data Fig. 5A were made using BioRender.

### Adoptive BMDC Transfer

C57BL/6 mice originally purchased from Charles River were bred and maintained at the Cross Cancer Institute vivarium. Mixed groups of male and female littermates 10-30 weeks old were used for experiments. All animal work was approved by the Cross Cancer Institute’s Animal Care Committee.

On day 1, 2×10^5^ CIN MC38 cells were injected orthotopically into the descending colon wall, as performed previously, using a flexible needle (Hamilton) inserted through the working channel of a Wolfe endoscope and visualized using the ColoView imaging system^53^.

Fresh BMDCs were prepared as described above. Following collection of non-adherent cells, BMDCs were CFDA stained at a ratio of 1×10^7^ BMDCs/1µM CFDA (Thermo Fisher Scientific) for 10 minutes at room temperature. BMDCs were then resuspended in RPMI (10% FBS, 1% Penicillin/Streptomycin, 1% HEPES, 50µM β-Mercaptoethanol) and allowed to recover for 30 minutes at 37°C. Following 2 PBS washes, BMDCs were resuspended in RPMI at 2×10^6^ cells/mL and stimulated using 2µg DNA and 0.125µL/mL Lipofectamine 2000 (Invitrogen) per 2×10^6^ cells for 30 minutes at 37°C as per manufacturer instructions. After 2 PBS washes, 2×10^6^ BMDCs were injected intraperitoneally into tumor bearing mice on days 1, 6, and 12.

On day 15, tumors, mesenteric lymph nodes, and spleens were collected, minced, and digested shaking for 30 minutes at 37°C in enzyme cocktail (RPMI, 10µg/mL DNaseI (Thermo Scientific), and 1mg/mL Collagenase IV (Sigma Aldrich)). Tissue was then vigorously pipetted to dissociate and filtered through a 100µm (tumor) or 40µm (mesenteric lymph node or spleen) strainer before ACK buffer (150mM Ammonium Chloride, 10mM Potassium Bicarbonate, 0.1mM EDTA, pH 7.4) removal of red blood cells (spleen only) and washing. Cells were restimulated for 4 hours with 0.5µg/mL PMA, 50µg/mL Ionomycin, and 2 hours with 2µM Monensin before flow cytometry staining using the antibodies in Supplemental Table 2, the Zombie Aqua Fixable Viability Kit (BioLegend), and the FOXP3/Transcription

Factor Staining Kit (eBioscience). Stained samples were run on the CytoFlex S flow cytometer (Beckman Coulter). Analysis was performed in FlowJo.

## Acknowledgements

The authors thank Dan McGinn, Cheryl Santos, Daming Li, Sudip Subedi, Jessica Hamilton, Dr. Xuejun Sun, Dr. Anne Galloway, and Dr. Lei Li as well as the Advanced Cell Exploration core at the University of Alberta for technical support. Patient samples for organoid development were provided by Dr. Daniel Schiller.

This project was supported by funding from the Canadian Institutes of Health Research (grant 407882), the Natural Sciences and Engineering Research Council of Canada (grant RGPIN-2016-05152), and the University Hospital Foundation (K. Baker).

## Supplemental Figures

**Extended Data Figure 1.**
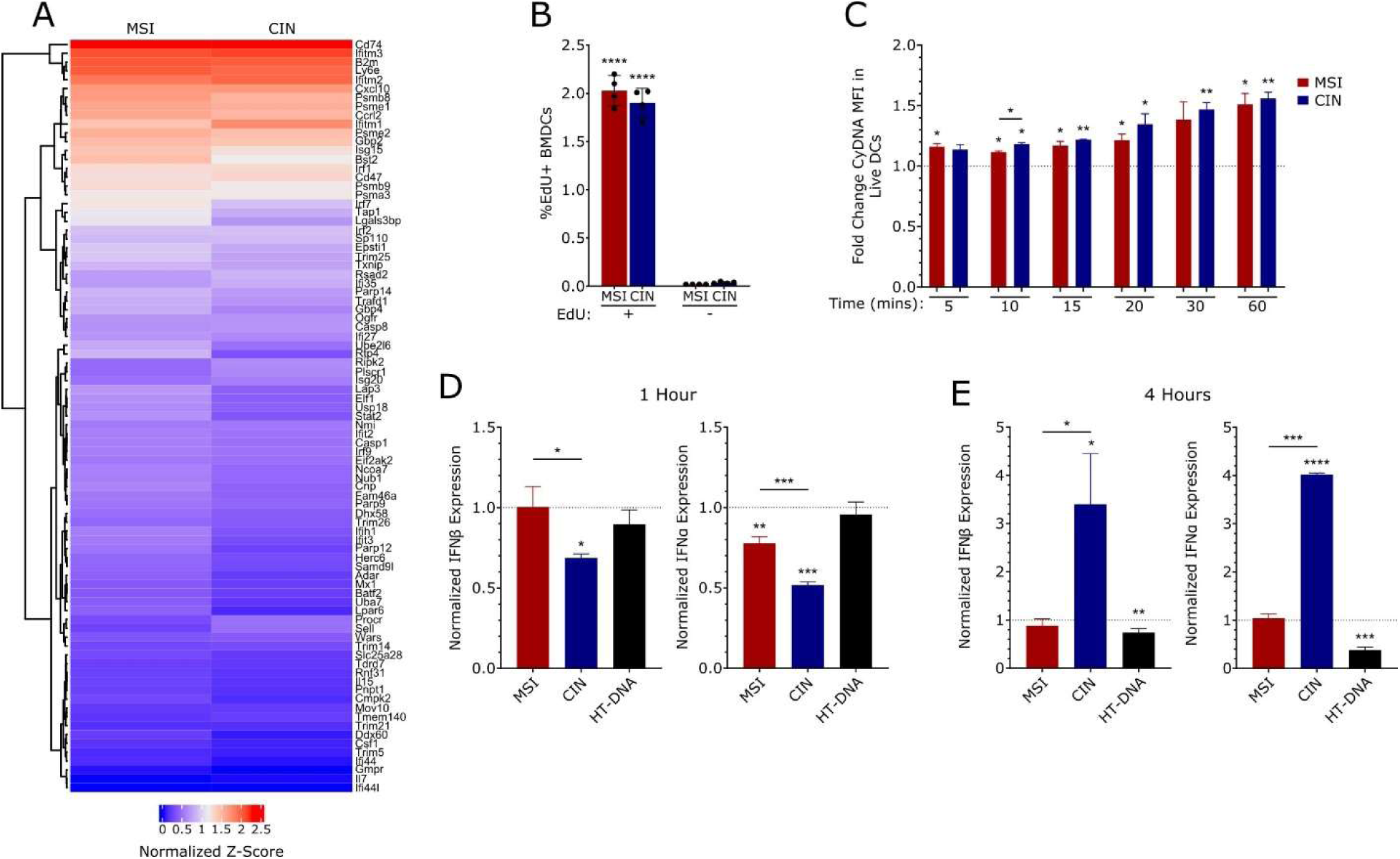
cyDNA from MSI cells leads to increased STING activation. (A) Z-score heat map of expression of the Type I IFN gene set in DCs from orthotopic MSI or CIN tumors. Cells were analyzed by single cell RNA sequencing. (B) BMDCs were co-cultured for 4 hours at a 1:1 ratio with MSI or CIN CRC cells stained with EdU before examination of CRC DNA uptake by BMDCs using flow cytometry. n = 3. (C) BMDCs were stimulated with 50ng PicoGreen stained cyDNA for the indicated times before analysis by flow cytometry to evaluate cyDNA uptake. Data shown as fold change to lipofectamine vehicle control. n = 2. (D, E) 500ng cyDNA isolated from MSI and CIN cells was used to stimulate BMDCs for 15 minutes before washing. BMDCs were then incubated for a total of 1 (D) or 4 (E) hours before RNA isolation and qPCR to examine STING activation. n = 3. Representative replicates shown for all panels. Statistical significance was determined by unpaired T-tests. Significance to lipofectamine vehicle control is shown over the sample bar, * = p < 0.05, ** = p < 0.01, *** = p < 0.001, **** = p < 0.0001.

**Extended Data Figure 2.**
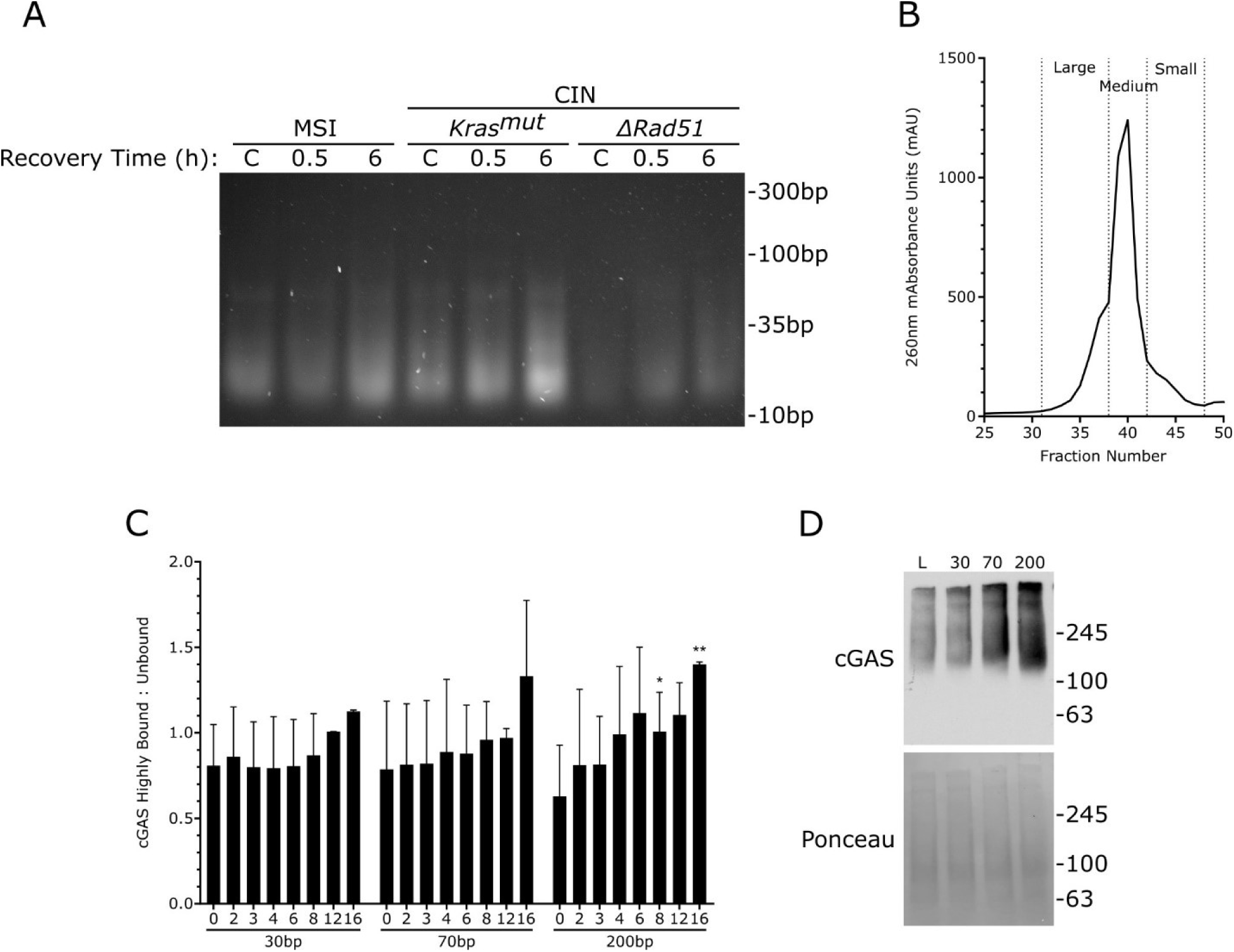
Increased cyDNA size leads to improved STING activation in DCs. (A) cyDNA isolated after 10 Gy of IR and the indicated recovery time was run on an 8% DNA PAGE gel and visualized. C indicates unirradiated cyDNA. n =2. (B) Fractionation demarcations for cyDNA isolated from MSI cells that was separated by FPLC and used for BMDC stimulation. n =3. (C) Quantification of the ratio of highly bound cGAS (defined as DNA remaining in the well) to unbound DNA oligos evaluated by EMSA (see Fig. 2F). Quantifications were performed with ImageJ. Statistical significance shown is in comparison to 30bp. n = 2. (D) BMDCs were stimulated for 40 minutes with 500ng of oligos of varying size before isolation of protein under non-denaturing conditions to examine cGAS oligomerization. n = 3. Statistical significance was determined by unpaired T-test, * = p < 0.05, ** = p < 0.01.

**Extended Data Figure 3.**
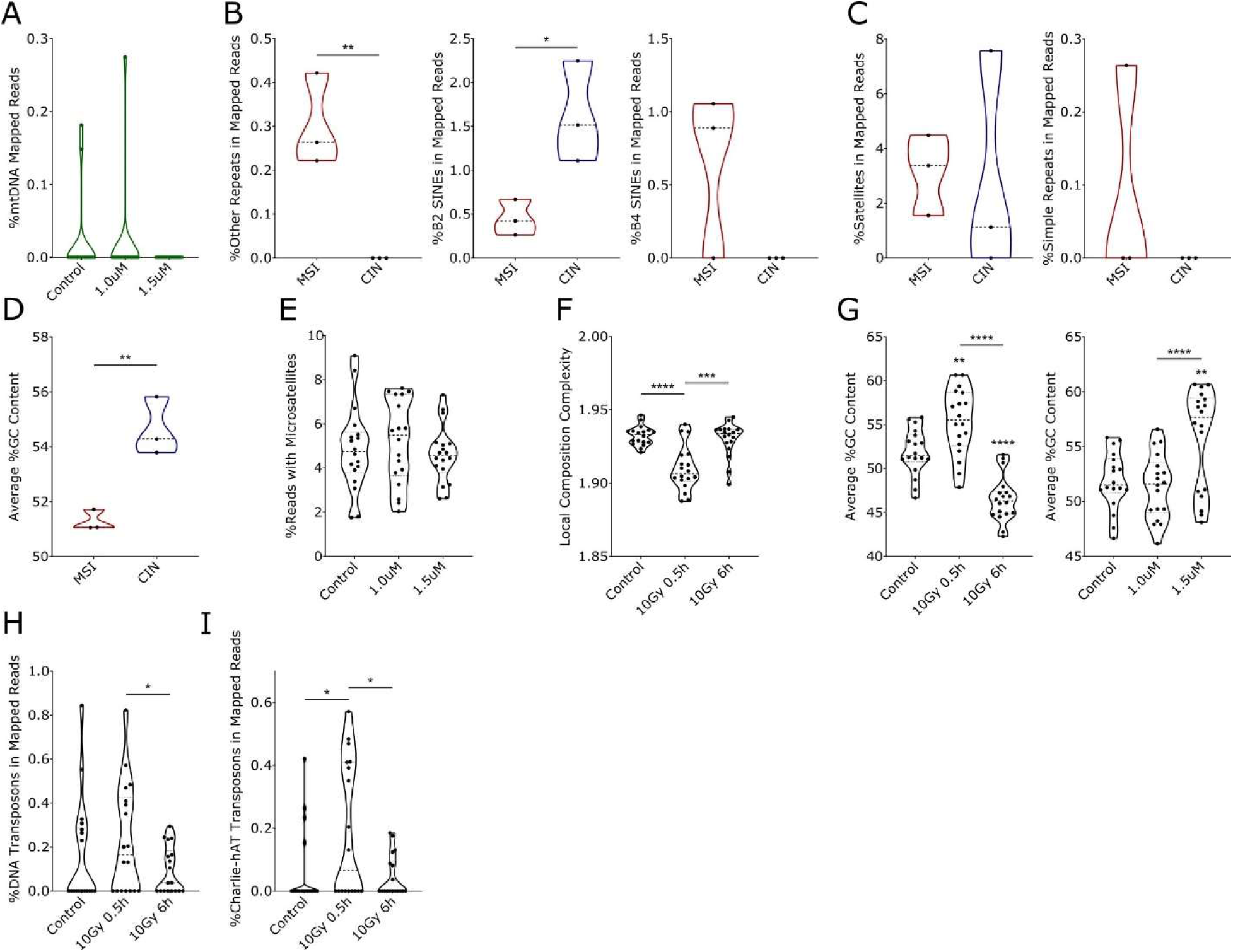
cyDNA differs in sequence depending on endogenous or treatment induced genetic instability. (A) mtDNA content in sequencing of cyDNA isolated following 5-FU treatment at the indicated dose for 24 hours. (B, C) Percent of mapped reads in RepeatMasker regions of the indicated repeat type in MSI and CIN cyDNA. (D) Percent GC of mapped and unmapped cyDNA sequencing reads in MSI vs CIN cells. (E) Percent of mapped and unmapped reads containing >5 2-6bp microsatellite repeats in cyDNA isolated following 5-FU treatment for 24 hours. (F) Local composition complexity of mapped and unmapped cyDNA sequencing reads of cyDNA from IR treated cells. (G) Percent GC of mapped and unmapped reads following IR (left) or 5-FU treatment (right). (H, I) Percent of mapped reads in RepeatMasker-annotated DNA transposon (H) and Charlie-hAT DNA transposon (I) regions in cyDNA isolated after IR treatment. All sequencing done with 3 biological replicates. Statistical significance was determined by paired (F, G, H, I) or unpaired (B, D) T-tests. Significance to untreated control is indicated over the sample (G), * = p < 0.05, ** = p < 0.01, *** = p < 0.001, **** = p < 0.0001.

**Extended Data Figure 4.**
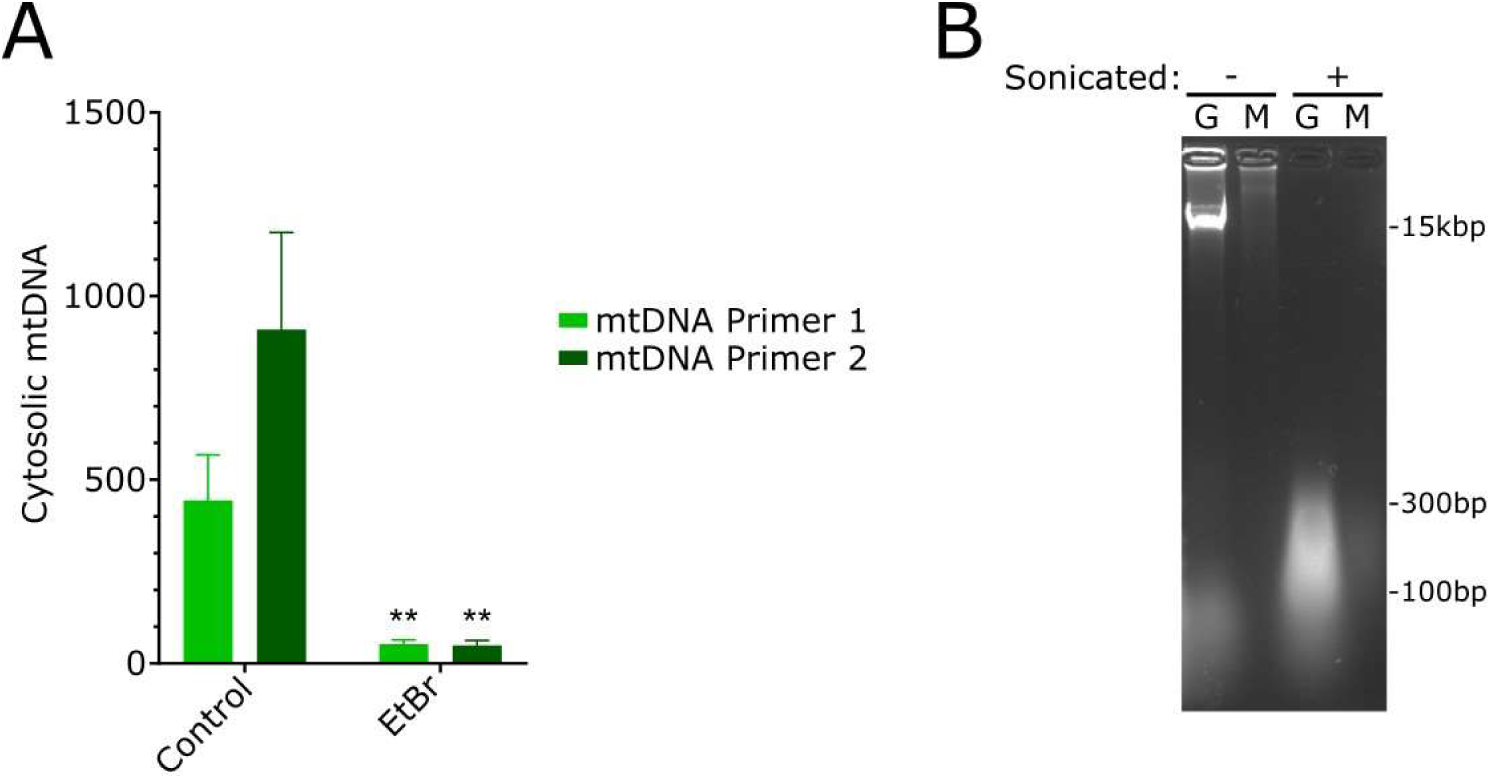
Confirmation of mtDNA depletion and equal mtDNA/genomic DNA size. (A) MSI cells were treated with EtBr for 7 days to deplete mtDNA before cyDNA isolation and determination of its mtDNA content by qPCR. Statistical significance was determined by unpaired T-tests, ** = p < 0.01. (B) mtDNA (M) and genomic DNA (G) were sonicated to equal size for stimulation of BMDCs (See Fig. 5C) and run on a 2% agarose gel.

**Extended Data Figure 5.**
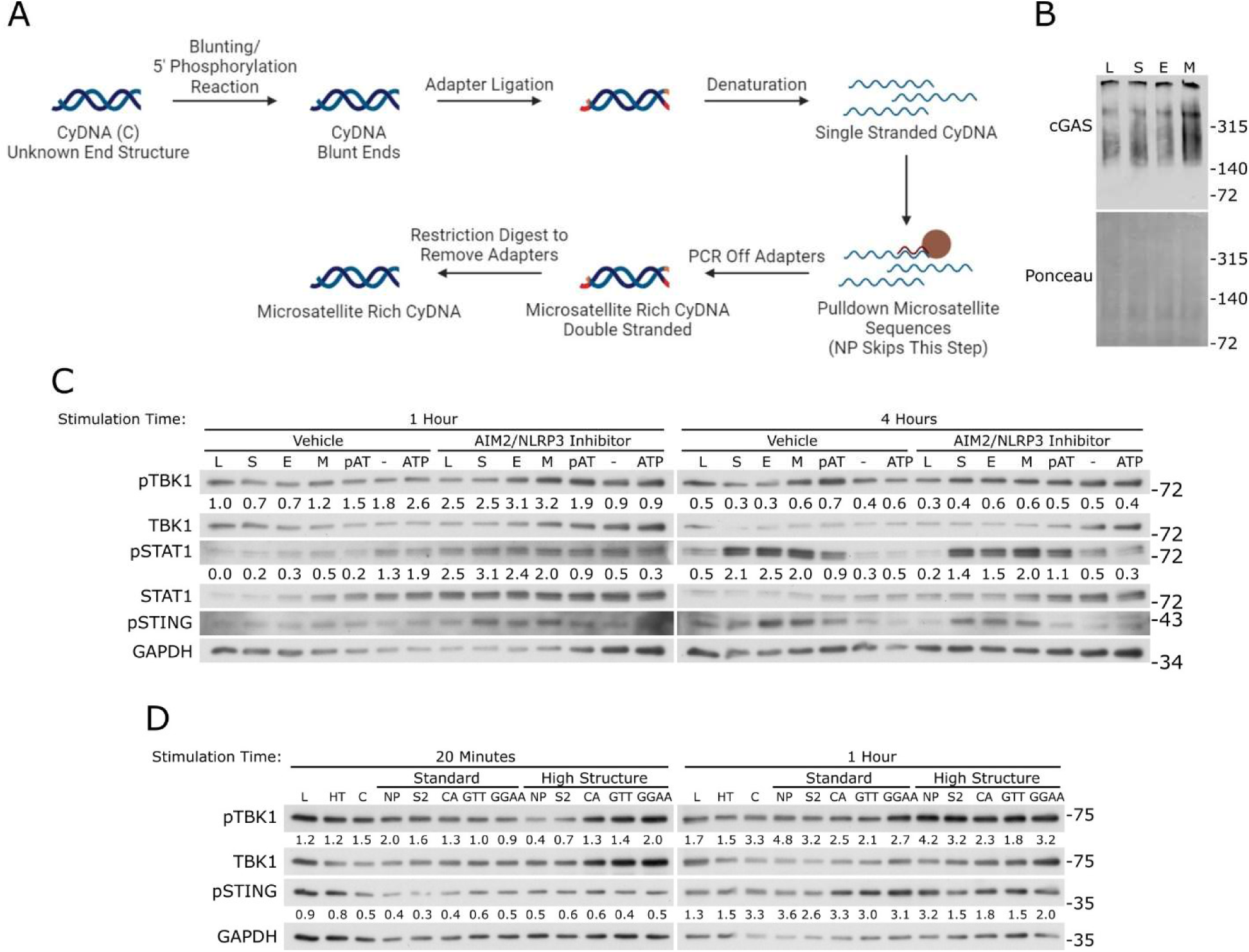
Microsatellites induce higher cGAS activation. (A) Schematic of microsatellite rich cyDNA pulldown protocol (See Fig. 6 C, D). (B) BMDCs were stimulated with oligos containing scrambled or microsatellite sequences or for 40 minutes before isolation and analysis of protein under non-denaturing conditions to examine cGAS oligomerization. L indicates lipofectamine vehicle control, S indicates scrambled oligo, E and M indicate edge and middle microsatellite oligos respectively. n = 3. (C) BMDCs were pre-treated for 30 minutes with the AIM2 and NLRP3 inhibitor IN-3 before stimulation with oligos containing scrambled or microsatellite sequences (oligo set #2) and protein isolation for western blot analysis. pAT refers to the AIM2 stimulus poly(dA:dT) (2µg), - indicates the media vehicle control for ATP stimulation, ATP was used at 2mM to stimulate NLRP3. n = 2. (D) Microsatellites were pulled down from MSI cyDNA as in A and Fig. 6C-D before re-annealing to induce highly structured DNA. BMDCs were stimulated with this DNA before isolation of protein and western blot analysis. C indicates unmanipulated MSI cyDNA, NP indicates cyDNA that was excluded from the pulldown step, S2 indicates pulldown with scrambled sequence, CA, GTT, and GGAA indicate microsatellites pulled down from MSI cyDNA. n = 2. Quantifications were performed in ImageJ and normalized to the GAPDH loading control.

**Extended Data Figure 6.**
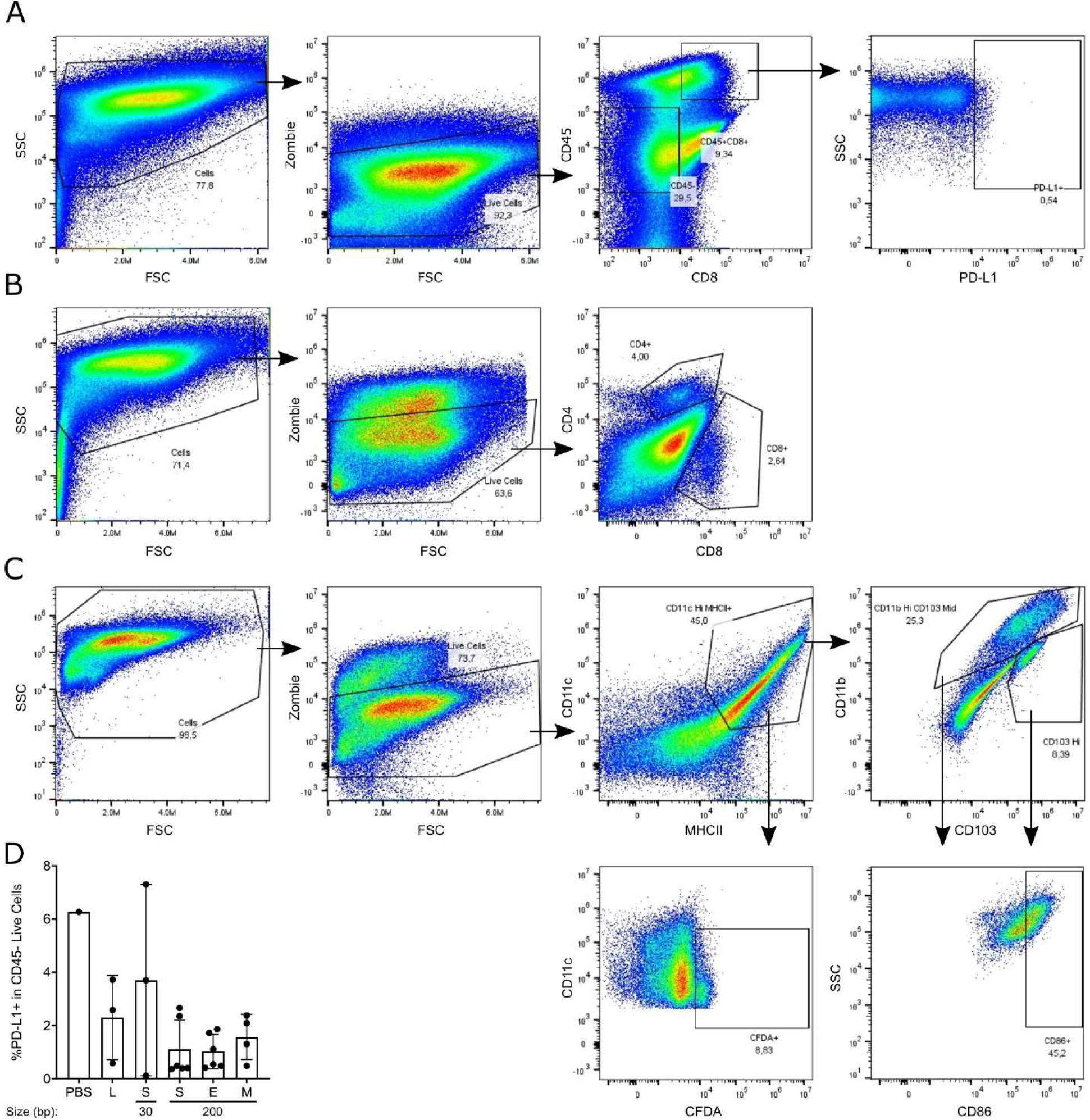
Treatment with microsatellite stimulated DCs does not increase PD-L1 levels on orthotopic CRC tumor cells. (A-C) Flow gating system for data in Fig. 7C left and Extended Data Fig. 6D (A), Fig. 7C Middle and Right (B), and Fig. 7D (C). (D) PD-L1 levels on CD45-tumor cells were evaluated by flow cytometry in orthotopic CIN tumors after 3 intraperitoneal injections with BMDCs that had been stimulated with oligos of the indicated length and containing scramble or microsatellite sequences (see Fig. 7). n = 3.

## Citations.

1. Xi, Y. & Xu, P. Global colorectal cancer burden in 2020 and projections to 2040. Transl. Oncol. 14, 101174 (2021).

2. Kuipers, E. J. et al. Colorectal cancer. Nat. Publ. Gr. 1, 1–25 (2015).

3. Dekker, E., Tanis, P. J., Vleugels, J. L. A., Kasi, P. M. & Wallace, M. B. Colorectal cancer. Lancet 394, 1467–1480 (2019).

4. Tennstedt, P., et al. RAD51 overexpression is a negative prognostic marker for colorectal adenocarcinoma. Int. J. Cancer 132, 2118–2126 (2013).

5. Kloor, M. & Von Knebel Doeberitz, M. The immune biology of microsatellite-unstable cancer. Trends in Cancer 2, 121–133 (2016).

6. Gelsomino, F., Barbolini, M., Spallanzani, A., Pugliese, G. & Cascinu, S. The evolving role of microsatellite instability in colorectal cancer: A review. Cancer Treat. Rev. 51, 19–26 (2016).

7. Llosa, N. J., et al. The vigorous immune microenvironment of microsatellite instable colon cancer is balanced by multiple counter-inhibitory checkpoints. Cancer Discov. 5, 43–51 (2015).

8. Kim, S. R., et al. Mismatch repair deficiency and prognostic significance in patients with low-risk endometrioid endometrial cancers. Int. J. Gynecol. Cancer 30, 783–788 (2020).

9. Mowat, C., Mosley, S. R., Namdar, A., Schiller, D. & Baker, K. Anti-tumor immunity in mismatch repair-deficient colorectal cancers requires type I IFN–driven CCL5 and CXCL10. J. Exp. Med. 218, (2021).

10. Schlee, M. & Hartmann, G. Discriminating self from non-self in nucleic acid sensing. Nat. Rev. Immunol. 16, 566–580 (2016).

11. Li, X., et al. Cyclic GMP-AMP Synthase Is Activated by Double-Stranded DNA-Induced Oligomerization. Immunity 39, 1019–1031 (2013).

12. Ng, K. W., Marshall, E. A., Bell, J. C. & Lam, W. L. cGAS–STING and Cancer: Dichotomous Roles in Tumor Immunity and Development. Trends Immunol. 39, 44–54 (2018).

13. Storozynsky, Q. & Hitt, M. M. The impact of radiation-induced dna damage on cgas-sting-mediated immune responses to cancer. Int. J. Mol. Sci. 21, 1–22 (2020).

14. Chabanon, R. M., et al. Targeting the DNA damage response in immuno-oncology: developments and opportunities. Nat. Rev. Cancer 21, (2021).

15. Zhu, Q. et al. Cutting Edge: STING Mediates Protection against Colorectal Tumorigenesis by Governing the Magnitude of Intestinal Inflammation. J. Immunol. 193, 4779–4782 (2014).

16. Deng, L., et al. STING-dependent cytosolic DNA sensing promotes radiation-induced type I interferon-dependent antitumor immunity in immunogenic tumors. Immunity 41, 843–852 (2014).

17. Flood, B. A., Higgs, E. F., Li, S., Luke, J. J. & Gajewski, T. F. STING pathway agonism as a cancer therapeutic. Immunol. Rev. 290, 24–38 (2019).

18. Schadt, L., et al. Cancer-Cell-Intrinsic cGAS Expression Mediates Tumor Immunogenicity. Cell Rep. 29, 1236–1248.e7 (2019).

19. Vanpouille-Box, C., Demaria, S., Formenti, S. C. & Galluzzi, L. Cytosolic DNA Sensing in Organismal Tumor Control. Cancer Cell 34, 361–378 (2018).

20. Rivera Vargas, T., Benoit-Lizon, I. & Apetoh, L. Rationale for stimulator of interferon genes– targeted cancer immunotherapy. Eur. J. Cancer 75, 86–97 (2017).

21. Andreeva, L., et al. CGAS senses long and HMGB/TFAM-bound U-turn DNA by forming protein-DNA ladders. Nature 549, 394–398 (2017).

22. Wiser, C., Kim, B., Vincent, J. & Ascano, M. Small molecule inhibition of human cGAS reduces total cGAMP output and cytokine expression in cells. Sci. Rep. 10, 1–11 (2020).

23. Longley, D. B., Harkin, D. P. & Johnston, P. G. 5-FLUOROURACIL : MECHANISMS OF ACTION AND CLINICAL STRATEGIES. 3, 330–338 (2003).

24. West, A. P., et al. Mitochondrial DNA stress primes the antiviral innate immune response. Nature 520, 553–557 (2015).

25. Yu, C. H., et al. TDP-43 Triggers Mitochondrial DNA Release via mPTP to Activate cGAS/STING in ALS. Cell 183, 636–649.e18 (2020).

26. De Cecco, M., et al. L1 drives IFN in senescent cells and promotes age-associated inflammation. Nature 566, 73–78 (2019).

27. Lindholm, H. T., Chen, R. & De Carvalho, D. D. Endogenous retroelements as alarms for disruptions to cellular homeostasis. Trends in Cancer 9, 55–68 (2023).

28. Smit, A., Hubley, R. & Green, P. RepeatMasker Open-4.0. http://www.repeatmasker.org.

29. Lo, J., Jonika, M. M. & Blackmon, H. micRocounter: Microsatellite characterization in genome assemblies. G3 Genes, Genomes, Genet. 9, 3101–3104 (2019).

30. Bourque, G., et al. Ten things you should know about transposable elements 06 Biological Sciences 0604 Genetics. Genome Biol. 19, 1–12 (2018).

31. Arensburger, P., et al. Phylogenetic and functional characterization of the hAT transposon superfamily. Genetics 188, 45–57 (2011).

32. Riley, J. S. & Tait, S. W. Mitochondrial DNA in inflammation and immunity . EMBO Rep. 21, 1–17 (2020).

33. Warren, E. B., Aicher, A. E., Fessel, J. P. & Konradi, C. Mitochondrial DNA depletion by ethidium bromide decreases neuronal mitochondrial creatine kinase: Implications for striatal energy metabolism. PLoS One 12, 1–22 (2017).

34. Xian, H., et al. Oxidized DNA fragments exit mitochondria via mPTP- and VDAC-dependent channels to activate NLRP3 inflammasome and interferon signaling. Immunity 55, 1370–1385.e8 (2022).

35. Hao, Z., et al. N6-Deoxyadenosine Methylation in Mammalian Mitochondrial DNA. Mol. Cell 78, 382–395.e8 (2020).

36. Zhong, F., Liang, S. & Zhong, Z. Emerging Role of Mitochondrial DNA as a Major Driver of Inflammation and Disease Progression. Trends Immunol. 40, 1120–1133 (2019).

37. Jiao, Y., et al. Discovery of a novel and potent inhibitor with differential species-specific effects against NLRP3 and AIM2 inflammasome-dependent pyroptosis. Eur. J. Med. Chem. 232, (2022).

38. Bagshaw, A. T. M. Functional mechanisms of microsatellite DNA in eukaryotic genomes. Genome Biol. Evol. 9, 2428–2443 (2017).

39. Le, D. T., et al. PD-1 Blockade in Tumors with Mismatch-Repair Deficiency. N. Engl. J. Med. 372, 2509–2520 (2015).

40. Wculek, S. K., et al. Dendritic cells in cancer immunology and immunotherapy. Nat. Rev. Immunol. 20, 7–24 (2020).

41. Guan, J., et al. MLH1 Deficiency-Triggered DNA Hyperexcision by Exonuclease 1 Activates the cGAS-STING Pathway. Cancer Cell 39, 109–121.e5 (2021).

42. Wang, J., et al. STING licensing of type I dendritic cells potentiates antitumor immunity. Sci. Immunol. 9, eadj3945 (2024).

43. Xie, W., et al. Human cGAS catalytic domain has an additional DNA-binding interface that enhances enzymatic activity and liquid-phase condensation. Proc. Natl. Acad. Sci. U. S. A. 116, 11946–11955 (2019).

44. Herzner, A. M., et al. Sequence-specific activation of the DNA sensor cGAS by Y-form DNA structures as found in primary HIV-1 cDNA. Nat. Immunol. 16, 1025–1033 (2015).

45. Gentili, M., Lahaye, X., Nadalin, F., Fachinetti, D. & Manel, N. The N-Terminal Domain of cGAS Determines Preferential Association with Centromeric DNA and Innate Immune Activation in the Nucleus Article The N-Terminal Domain of cGAS Determines Preferential Association with Centromeric DNA and Innate Immune Activation. 2377–2393 (2019) doi:10.1016/j.celrep.2019.01.105.

46. Balzarolo, M., et al. M6A methylation potentiates cytosolic dsDNA recognition in a sequence-specific manner. Open Biol. 11, (2021).

47. MacDonald, K. M., et al. Antecedent chromatin organization determines cGAS recruitment to ruptured micronuclei. Nat. Commun. 14, 1–15 (2023).

48. Baker, K., et al. Neonatal Fc receptor for IgG (FcRn) regulates cross-presentation of IgG immune complexes by CD8-CD11b+ dendritic cells. Proc. Natl. Acad. Sci. U. S. A. 108, 9927–9932 (2011).

49. Sato, T., et al. Long-term expansion of epithelial organoids from human colon, adenoma, adenocarcinoma, and Barrett’s epithelium. Gastroenterology 141, 1762–1772 (2011).

50. Miyoshi, H. & Stappenbeck, T. S. In vitro expansion and genetic modification of gastrointestinal stem cells in spheroid culture. Nat. Protoc. 8, 2471–2482 (2013).

51. Moffat, J., et al. A Lentiviral RNAi Library for Human and Mouse Genes Applied to an Arrayed Viral High-Content Screen. Cell 124, 1283–1298 (2006).

52. Koo, B. K., et al. Controlled gene expression in primary Lgr5 organoid cultures. Nat. Methods 9, 81–83 (2012).

53. Roper, J., et al. Colonoscopy-based colorectal cancer modeling in mice with CRISPR-Cas9 genome editing and organoid transplantation. Nat. Protoc. 13, 217–234 (2018).

54. Mosley, S. R. & Baker, K. Isolation of endogenous cytosolic DNA from cultured cells. STAR Protoc. 3, 101165 (2022).

55. Subramanian, A., et al. Gene set enrichment analysis: A knowledge-based approach for interpreting genome-wide expression profiles. Proc. Natl. Acad. Sci. U. S. A. 102, 15545–15550 (2005).

56. Cock, P. J. A., et al. Biopython: Freely available Python tools for computational molecular biology and bioinformatics. Bioinformatics 25, 1422–1423 (2009).

57. Andrews, S. Fast QC: A quality control tool for high throughput sequence data [online]. Available online at: http://www.bioinformatics.babraham.ac.uk/projects/fastqc/. (2010).

58. Morgan, M., Pages, H., Obenchain, V. & Hayden, N. Rsamtools: Binary alignment (BAM), FASTA, variant call (BCF), and tabix file import. R package version 2.10.0. https://bioconductor.org/packages/Rsamtools (2021).

59. Lawrence, M., Gentleman, R. & Carey, V. rtracklayer: An R package for interfacing with genome browsers. Bioinformatics 25, 1841–1842 (2009).

60. Lawrence, M. et al. Software for Computing and Annotating Genomic Ranges. PLoS Comput. Biol. 9, 1–10 (2013).

61. Yue, F., et al. A comparative encyclopedia of DNA elements in the mouse genome. Nature 515, 355–364 (2014).

62. Bushnell, B., Rood, J. & Singer, E. BBMerge – Accurate paired shotgun read merging via overlap. PLoS One 12, 1–15 (2017).

63. St. John, J. & Quinn, T. W. Rapid capture of DNA targets. Biotechniques 44, 259–264 (2008).

