## Supplemental Tables for "Mechanisms of genomic instability dictate cytosolic DNA composition and dendritic cell mediated anti-tumor immunity"

**Supplemental Table 1. DNA oligo sequences.**

| Oligo Name | Relevant Figures | Sequence |
|---|---|---|
| 30bp Oligo 1 Fwd | Fig. 2B | TCTTTCAGAGGCCCGCGTTTACGACTTCAT |
| 30bp Oligo 1 Rev | Fig. 2B | ATGAAGTCGTAAACGCGGGCCTCTGAAAGA |
| 30bp Oligo 2 Fwd | Fig. 2B, 2D-F, 7 Extended Data Fig. 2C-D, 6 | AATATACCGTGTTCAGCCGGGTAGCTTATT |
| 30bp Oligo 2 Rev | Fig. 2B, 2D-F, 7 Extended Data Fig. 2C-D, 6 | AATAAGCTACCCGGCTGAACACGGTATATT |
| 30bp Oligo 3 Fwd | Fig. 2B | GCGGCTGTGATGTTTTGCCATCGAGAAGAT |
| 30bp Oligo 3 Rev | Fig. 2B | ATCTTCTCGATGGCAAAACATCACAGCCGC |
| 50bp Oligo 1 Fwd | Fig. 2B | CCTCACGTCGAGTTTCAGTGTCGGCATATGCTATCCGTAGCAAAATTCCC |
| 50bp Oligo 1 Rev | Fig. 2B | GGGAATTTTGCTACGGATAGCATATGCCGACACTGAAACTCGACGTGAGG |
| 50bp Oligo 2 Fwd | Fig. 2B | GTCATGAATCAATCGCACTGAGTGAATTCGTGAAAGAAATGAGGTAGCTT |
| 50bp Oligo 2 Rev | Fig. 2B | AAGCTACCTCATTTCTTTCACGAATTCACTCAGTGCGATTGATTCATGAC |
| 50bp Oligo 3 Fwd | Fig. 2B | GCGATCGTCTTATCTGCACGCGTGCAAACTCCAAGCTACCATTGCGTGTA |
| 50bp Oligo 3 Rev | Fig. 2B | TACACGCAATGGTAGCTTGGAGTTTGCACGCGTGCAGATAAGACGATCGC |
| 70bp Oligo 1 Fwd | Fig. 2B | CTCCGGCGCCAAAGAGTATTGGGTCCTATCCACTGGTCAGCAAACTGTGCCATATGAAGCAAGTAACACG |
| 70bp Oligo 1 Rev | Fig. 2B | CGTGTTACTTGCTTCATATGGCACAGTTTGCTGACCAGTGGATAGGACCCAATACTCTTTGGCGCCGGAG |
| 70bp Oligo 2 Fwd | Fig. 2B, 2D-F, Extended Data Fig. 2C-D | GGCAAGCAGGCCTTCTCATGCGTATATCAACGTGTCCGTTTGAGATATCAGTGTTCTCTTTCCCGGTTAA |
| 70bp Oligo 2 Rev | Fig. 2B, 2D-F, Extended Data Fig. 2C-D | TTAACCGGGAAAGAGAACACTGATATCTCAAACGGACACGTTGATATACGCATGAGAAGGCCTGCTTGCC |
| 70bp Oligo 3 Fwd | Fig. 2B | GGCACTCAGATGCATCGAACATCGGCGGATAGCTCCCGGTAAGAAGTAAGACATGTACCCAAGTTGTAGT |
| 70bp Oligo 3 Rev | Fig. 2B | ACTACAACTTGGGTACATGTCTTACTTCTTACCGGGAGCTATCCGCCGATGTTCGATGCATCTGAGTGCC |
| 200bp Oligo Fwd | Fig. 2D-F, Extended Data Fig. 2C-D | TTAATAACATCGAGCATTAGGGAGCCGCTAGTTCGATTGTATTAATAACAAGGACTGTCCCTATTAACCTTGCCGGATCTGAAGCGACACCGTGTTACTTGATGCCACAAGAGTAACACGAACTTAATTATCGCCGTCGGGAGCGTTGCCTGTCGTCTTCTAGCCAATGTATACCATCAAGGTGAACCGCTTAGTTAGGA |
| 200bp Oligo Rev | Fig. 2D-F, Extended Data Fig. 2C-D | TCCTAACTAAGCGGTTCACCTTGATGGTATACATTGGCTAGAAGACGACAGGCAACGCTCCCGACGGCGATAATTAAGTTCGTGTTACTCTTGTGGCATCAAGTAACACGGTGTCGCTTCAGATCCGGCAAGGTTAATAGGGACAGTCCTTGTTATTAATACAATCGAACTAGCGGCTCCCTAATGCTCGATGTTATTAA |

| | | |
|---|---|---|
| Scramble 1 Fwd | Fig. 6A-B | TTAATAACATCGAGCATTAGGGAGCCGCTAGTTCGATTGTATTAATAACAAGGACTGTCCCTATTAACCTTGCCGGATCTGAAGCGACACCGTGTTACTTGATGCCACAAGAGTAACACGAACTTAATTATCGCCGTCGGGAGCGTTGCCTGTCGTCTTCTAGCCAATGTATACCATCAAGGTGAACCGCTTAGTTAGGA |
| Scramble 1 Rev | Fig. 6A-B | TCCTAACTAAGCGGTTCACCTTGATGGTATACATTGGCTAGAAGACGACAGGCAACGCTCCCGACGGCGATAATTAAGTTCGTGTTACTCTTGTGGCATCAAGTAACACGGTGTCGCTTCAGATCCGGCAAGGTTAATAGGGACAGTCCTTGTTATTAATACAATCGAACTAGCGGCTCCCTAATGCTCGATGTTATTAA |
| Scramble 2 Fwd | Fig. 6A-B, 6E-G, 7, Extended Data Fig. 5B-C, 6 | ATGTCCTGGCACCCTGAAAAATGCTGATGAGGTAATGGTCTACTAGCTTCCTATTTTCGCGGTCGAGGGTTGGCATGTAAAATCCAGTGATTGACATAGCACGCACACACTGTCCCTACATGGTTCGCACAAGTGTGACCAGACATACAGCGTCAAACGGGAATCAACTAAGGACTAGAGGCCCACTACCTATAGAGATG |
| Scramble 2 Rev | Fig. 6A-B, 6E-G, 7, Extended Data Fig. 5B-C, 6 | CATCTCTATAGGTAGTGGGCCTCTAGTCCTTAGTTGATTCCCGTTTGACGCTGTATGTCTGGTCACACTTGTGCGAACCATGTAGGGACAGTGTGTGCGTGCTATGTCAATCACTGGATTTTACATGCCAACCCTCGACCGCGAAAATAGGAAGCTAGTAGACCATTACCTCATCAGCATTTTTCAGGGTGCCAGGACAT |
| Microsatellite Edge 1 Fwd | Fig. 6A-B | AGTTGCAGACGTTGGCTCAGGTCCAGGCTTATTGTAGCAAAGTCTACAAAGCAGTTCTAAGTGACCTTTATTGTGACTGGCTCTAAAGGAAGGCGAGAGCCTGGTCTACAAAGTGAGCTCCAAGACAGCCAGGGCTACACAGAGAAACCCTGTCTGGAAAAAACAAAAAACAAAAAACAACAACAACAACAACAAAAAAG |
| Microsatellite Edge 1 Rev | Fig. 6A-B | CTTTTTTGTTGTTGTTGTTGTTGTTTTTTGTTTTTTGTTTTTTCCAGACAGGGTTTCTCTGTGTAGCCCTGGCTGTCTTGGAGCTCACTTTGTAGACCAGGCTCTCGCCTTCCTTTAGAGCCAGTCACAATAAAGGTCACTTAGAACTGCTTTGTAGACTTTGCTACAATAAGCCTGGACCTGAGCCAACGTCTGCAACT |
| Microsatellite Edge 2 Fwd | Fig. 6A-B, 6E-G, 7, Extended Data Fig. 5B-C, 6 | AACAACAACAACAACAACAACAACAAAAAAAAAAAAAACTCTGCTTACCCTATGTAGTAACTTCTGCACCAGTGGAGGACAGAGGAAGATGTAGGGTGTGCTGTTCTAATATTCCCACTCTCCTTCTTCACCATATCTTACCAACAGAGAACATGGGGAAAATGGGCAGACATAGCTGTCTTTCTAAATGAATATTAGGCA |
| Microsatellite Edge 2 Rev | Fig. 6A-B, 6E-G, 7, Extended Data Fig. 5B-C, 6 | TGCCTAATATTCATTTAGAAAGACAGCTATGTCTGCCCATTTTCCCCATGTTCTCTGTTGGTAAGATATGGTGAAGAAGGAGAGTGGGAATATTAGAACAGCACACCCTACATCTTCCTCTGTCCTCCACTGGTGCAGAAGTTACTACATAGGGTAAGCAGAGTTTTTTTTTTTTTTGTTGTTGTTGTTGTTGTTGTTGTT |
| Microsatellite Middle 1 Fwd | Fig. 6A-B | TAAGCATCAAAACCAATTTCCTTACCAGGAGGCAATCATGCATATGGTACAAGGAGGAAACTAACAAAAACAAAAAGACCAAAAAACCCAAAAACAACAACAACAACAACAAAAACCTGTGCCTTATCCAATGCTATCCACCTTCATCCACTAACTGCATAATAGAATGCAAATGGTAAGCTAGATGCCTCATGATTGCT |
| Microsatellite Middle 1 Rev | Fig. 6A-B | AGCAATCATGAGGCATCTAGCTTACCATTTGCATTCTATTATGCAGTTAGTGGATGAAGGTGGATAGCATTGGATAAGGCACAGGTTTTTGTTGTTGTTGTTGTTGTTTTTGGGTTTTTTGGTCTTTTTGTTTTTGTTAGTTTCCTCCTTGTACCATATGCATGATTGCCTCCTGGTAAGGAAATTGGTTTTGATGCTTA |
| Microsatellite Middle 2 Fwd | Fig. 6A-B, 6E-G, 7, Extended Data Fig. 5B-C, 6 | TTGGTTATCATCACCAGGGCCCGGGAGAAACATTTTTTACTTCCTTTTCATTTCTTCTATCCTAGTCTGGTTCAGTTGCTGTGATTTAAAACAAAACAAAACAACAACAACAACAACAAAACAAAAAACAACTTGGGGA |

| | | |
|---|---|---|
| | | AGAAAGTGTTTCTTAGGCTCACACATCCTGATCATAGTCCATCAAGGAGGGAAGTCAGGGC |
| Microsatellite Middle 2 Rev | Fig. 6A-B, 6E-G, 7, Extended Data Fig. 5B-C, 6 | GCCCTGACTTCCCTCCTTGATGGACTATGATCAGGATGTGTGAGCCTAAGAAACACTTTCTTCCCCAAGTTGTTTTTTGTTTTGTTGTTGTTGTTGTTGTTTTGTTTTGTTTTAAATCACAGCAACTGAACCAGACTAGGATAGAAGAAATGAAAAGGAAGTAAAAAATGTTTCTCCCGGGCCCTGGTGATGATAACCAA |
| 10bp Scramble Probe (S2) | Fig. 6C-D, Extended Data Fig. 5D | Biotin-TACAGCGACG |
| 15bp Scramble Probe (S3) | Fig. 6C-D | Biotin-CAGCATTAGCGTAGA |
| $CA_5$ Probe | Fig. 6C-D, Extended Data Fig. 5D | Biotin-CACACACACA |
| $TC_5$ Probe | Fig. 6C-D | Biotin-TCTCTCTCTC |
| $TG_5$ Probe | Fig. 6C-D | Biotin-TGTGTGTGTG |
| $TGG_5$ Probe | Fig. 6C-D | Biotin-TGGTGGTGGTGGTGG |
| $GTT_5$ Probe | Fig. 6C-D, Extended Data Fig. 5D | Biotin-GTTGTTGTTGTTGTT |
| $GGAA_5$ Probe | Fig. 6C-D, Extended Data Fig. 5D | Biotin-GGAAGGAAGGAAGGAAGGAA |
| Pulldown Adapter 1 Fwd | Fig. 6C-D, Extended Data Fig. 5D | GTCTCCTCTGACTTCAACAGCGGATATC |
| Pulldown Adapter 1 Rev | Fig. 6C-D, Extended Data Fig. 5D | GATATCCGCTGTTGAAGTCAGAGGAGACC |
| Pulldown Adapter 2 Fwd | Fig. 6C-D, Extended Data Fig. 5D | GATATCACCACCCTGTTGCTGTAGCCAA |
| Pulldown Adapter 2 Rev | Fig. 6C-D, Extended Data Fig. 5D | TTTGGCTACAGCAACAGGGTGGTGATATC |

**Supplemental Table 2. Antibody list.**

| Target | Company | Catalog Number |
|---|---|---|
| IFNg-PE | Invitrogen | 12-7311-82 |
| CD69-$A^{700}$ | Invitrogen | 56-0691-82 |
| Ki67-PECy5 | Invitrogen | 15-5698-82 |
| pTBK1 | Cell Signaling Technology | 5483S |
| TBK1 | Cell Signaling Technology | 3504S |
| pSTAT1 | Invitrogen | 33-3400 |
| STAT1 | Cell Signaling Technology | 9172L |
| GAPDH | Invitrogen | MA5-15738 |
| pSTING | Cell Signaling Technology | 72971S |
| βActin | Cell Signaling Technology | 8457L |
| pNF-κB | Cell Signaling Technology | 3033S |
| cGAS | Cell Signaling Technology | 31659S |
| Anti-Rabbit IgG | Invitrogen | 26102 |
| MHCII-PECy5 | Invitrogen | 15-5321-82 |
| CD8-$A^{700}$ | Invitrogen | 56-0081-82 |
| CD45-PE | BioLegend | 103106 |
| CD86-APCCy7 | Invitrogen | 47-0862-82 |
| CD103-PE | Invitrogen | 12-1031-83 |
| CD11c-$A^{700}$ | BD Biosciences | 560583 |
| CD11b-PECy7 | Invitrogen | 25-0112-82 |
| PD-L1-$A^{750}$ | Bioss | Bs-4941R-A750 |
| Anti-Rabbit HRP Secondary | Jackson Immunoresearch | 111-035-144 |
| Anti-Mouse HRP Secondary | Jackson Immunoresearch | 115-035-146 |
| $A^{594}$ Secondary | Fisher Scientific | 111585045 |

**Supplemental Table 3. Primer list.**

| Primer Name | Sequence |
|---|---|
| GAPDH qPCR Fwd | CATGTTCCAGTATGACTCCA |
| GAPDH qPCR Rev | TGAAGACACCAGTAGACTCC |
| IFNβ qPCR Fwd | CGTGGGAGATGTCCTCAACT |
| IFNβ qPCR Rev | AGATCTCTGCTCGGACCACC |
| CXCL10 qPCR Fwd | CCAAGTGCTGCCGTCATTTTC |
| CXCL10 qPCR Rev | GGCTCGCAGGGATGATTTCAA |
| IRF7 qPCR Fwd | GGTGTGTCCCCAGGATCATT |
| IRF7 qPCR Fwd | GCTGCATAGGGTTCCTCGTAA |
| IFNα qPCR Fwd | GGATGTGACCTTCCTCAGACTC |
| IFNα qPCR Rev | ACCTTCTCCTGCGGGAATCCAA |
| mtDNA 1 qPCR Fwd | AACGGATCCACAGCCGTA |
| mtDNA 1 qPCR Rev | AGTCCTCGGGCCATGATT |
| mtDNA 2 qPCR Fwd | CAAACACTTATTACAACCCAAGAACA |
| mtDNA 2 qPCR Rev | TCATATTATGGCTATGGGTCAGG |
| Pulldown Adapter PCR Fwd | GTCTCCTCTGACTTCAACAGCG |
| Pulldown Adapter PCR Rev | ACCACCCTGTTGCTGTAGCCAA |
| Scramble 2 Oligo PCR Fwd | CACACTGTCCCTACATGGTTC |
| Scramble 2 Oligo PCR Rev | TTTGACGCTGTATGTCTGGTC |
| Microsatellite Edge 2 PCR Fwd | GATGTAGGGTGTGCTGTTCTAA |
| Microsatellite Edge 2 PCR Rev | CCCATGTTCTCTGTTGGTAAGA |
| Microsatellite Middle 2 PCR Fwd | GAAAGTGTTTCTTAGGCTCACAC |
| Microsatellite Middle 2 PCR Rev | CTTCCCTCCTTGATGGACTATG |
